# Interplay between Desmoglein2 and hypoxia controls metastasis in breast cancer

**DOI:** 10.1101/790519

**Authors:** Po-Hao Chang, Min-Che Chen, Ya-Ping Tsai, Grace Y.T. Tan, Pang-Hung Hsu, Yung-Ming Jeng, Yi-Fang Tsai, Muh-Hwa Yang, Wendy W. Hwang-Verslues

## Abstract

Metastasis is the major cause of cancer death. An increased level of circulating tumor cells (CTCs), metastatic cancer cells that have intravasated into the circulatory system, is particularly associated with colonization of distant organs and poor prognosis. However, the key factors required for tumor cell dissemination and colonization remain elusive. We found that high expression of Desmoglein2 (DSG2), a component of desmosome-mediated intercellular adhesion complexes, promoted tumor growth, increased the prevalence of CTC clusters and facilitated distant organ colonization. The dynamic regulation of DSG2 by hypoxia was key to this process as downregulation of DSG2 in hypoxic regions of primary tumors led to elevated epithelial-mesenchymal transition (EMT) gene expression, allowing cells to detach from the primary tumor and undergo intravasation. Subsequent derepression of DSG2 after intravasation and release of hypoxic stress was associated with an increased ability to colonize distant organs. This dynamic regulation of DSG2 was mediated by Hypoxia-Induced Factor1α (HIF1α). In contrast to its more widely observed function to promote expression of hypoxia-inducible genes, HIF1α repressed DSG2 by recruitment of the Polycomb Repressive Complex 2 components, EZH2 and SUZ12, to the DSG2 promoter in hypoxic cells. Consistent with our experimental data, DSG2 expression level correlated with poor prognosis and recurrence risk in breast cancer patients. Together, these results demonstrated the importance of DSG2 expression in metastasis and revealed a new mechanism by which hypoxia drives metastasis.

**Significance Statement:** During metastasis, hypoxia is a major force driving primary tumor cells to disseminate into the circulatory system. The key factors that promote circulating tumor cells (CTC) dissemination and allow them to successfully colonize distal sites are still incompletely known. We found that downregulation of DSG2 in hypoxic tumor allowed single tumor cell dissemination while DSG2 expressing tumors generated more CTC clusters. Re-induction of DSG2 expression in single CTCs may contribute to CTC survival and colonization in distant organs. These findings highlight the importance of DSG2 in breast cancer progression and metastasis.

## Introduction

Breast cancer is the most common cancer of women worldwide. With early detection and advances in therapeutic strategies, the 5-year relative survival rate for all stages combined is higher than 90% (according to the SEER database maintained by the U.S. National Cancer Institute). However, metastasis is now the major cause of breast cancer death (1) as the survival rate for women with metastasized breast cancers remains below 30%. Thus, identification of key factors in breast tumorigenesis and metastasis is important to identify therapeutic targets and strategies to improve prognosis.

Metastasis is a complex process involving tumor cell intrinsic alterations and extrinsic interaction with the microenvironment to select for highly aggressive cancer cells. Hypoxia, a key microenvironmental factor in solid tumors, activates hypoxic signaling to increase plasticity and promote epithelial-mesenchymal transition (EMT) to drive the first step of metastasis (2). High plasticity allows cancer cells to disseminate from the primary site and intravasate into the circulatory or lymphatic system. Most of these circulating tumor cells (CTCs) die in circulation and only a small fraction of CTCs are able to survive and eventually colonize distant organs (3, 4). Recent evidence has shown that CTC numbers can be used as an independent predictor for survival in patients with metastatic cancers (5–7). Further improvement in CTC detection methods led to identification of CTC clusters and the finding that clustered CTCs exhibit epithelial/mesenchymal hybrid (partial EMT) phenotype which allows them to move collectively (8). Collective movement makes these cancer cells more apoptosis-resistant, more capable of avoiding immune surveillance and, better able to colonize distant organs. Importantly, CTC clustering ability has been positively correlated with poor clinical outcome (9–12). While a few factors that promote CTC cluster formation have been identified (9, 13–15), the key mechanisms that allow CTC clusters to survive in the vascular system and allow them to more efficiently metastasize than unclustered CTCs remain elusive.

Cell adhesion proteins play critical roles in intercellular contacts and epithelial tissue dynamics. Deregulation of cell adhesion molecules contributes to tumor metastasis (16, 17). Among cell adhesion molecules, desmosomes are of particular interest for cancer biology. Desmosomes form patch-like adhesion structures that mark the intercellular midline and connect to the intermediate filament cytoskeleton to maintain cell-cell adhesion and tissue integrity (18). The desmosome is a protein complex containing two transmembrane proteins, Desmocollin (DSC1-DSC3) and Desmoglein (DSG1-DSG4), as well as adaptor proteins, Plakoglobin and Desmoplakin, that bind intermediate filaments (19). Among the human DSGs, DSG1 and DSG3 expression is mainly restricted to stratified squamous epithelia (20). DSG2 is the most ubiquitously expressed isoform, including mammary tissue, and is a key factor for cell aggregation and oncogenic function in lung and prostate cancers (20–23). However, whether DSG2 is involved in CTC clustering and metastasis remains unknown.

We found that tumor growth and colonization were promoted by DSG2 expression at both the primary site and distant organ. High DSG2 expression in the primary tumor was associated with increased prevalence of CTC clusters. However, HIF1αmediated suppression of DSG2 under hypoxia was required for cancer cell invasion and migration. Once in the vascular system, the cancer cells were released from hypoxic stress and DSG2 expression was de-repressed. This DSG2 reactivation was essential for CTCs to colonize distant organs. Consistent with these experimental observations, clinical data indicated that breast cancer patients whose tumors expressed DSG2 (DSG2-positive) had worse prognosis and higher recurrence risk than those with DSG2-negative tumors. Together, these results show that dynamic changes of DSG2 expression are required for breast tumorigenesis, CTC clustering, invasion and metastasis. Our data also identify regulatory mechanisms underlying DSG2 repression and de-repression during specific stages of breast cancer progression.

## Results

### High expression of DSG2 in breast cancer is associated with poor prognosis and high recurrence

To identify key factors driving CTC clustering and metastasis, we analyzed a transcriptome dataset for metastatic and non-metastatic breast cancers from the Gene Expression Omnibus (GEO) database using Gene Set Enrichment Analysis (GSEA). Forty-six genes associated with metastatic disease were enriched in metastatic breast cancers. Among these, DSG2 was the most highly expressed (Fig. 1A, 1B). Because formation of CTC clusters (groups of two or more aggregated CTCs) promotes metastasis and because cell-cell interaction is critical for CTC clustering (9), we also analyzed genes associated with cell-cell junctions and found DSG2 as the second most highly expressed gene among 56 genes enriched in the metastatic group (*SI Appendix*, Fig. S1A, S1B). Overlap of these two gene sets (metastasis and cell-cell junction) identified three genes (*SI Appendix*, Fig. S1C). Among these, DSG2 was the most highly upregulated in the metastasis and cell-cell junction gene sets that was also enriched in metastatic breast cancer patients.

**Figure 1.**
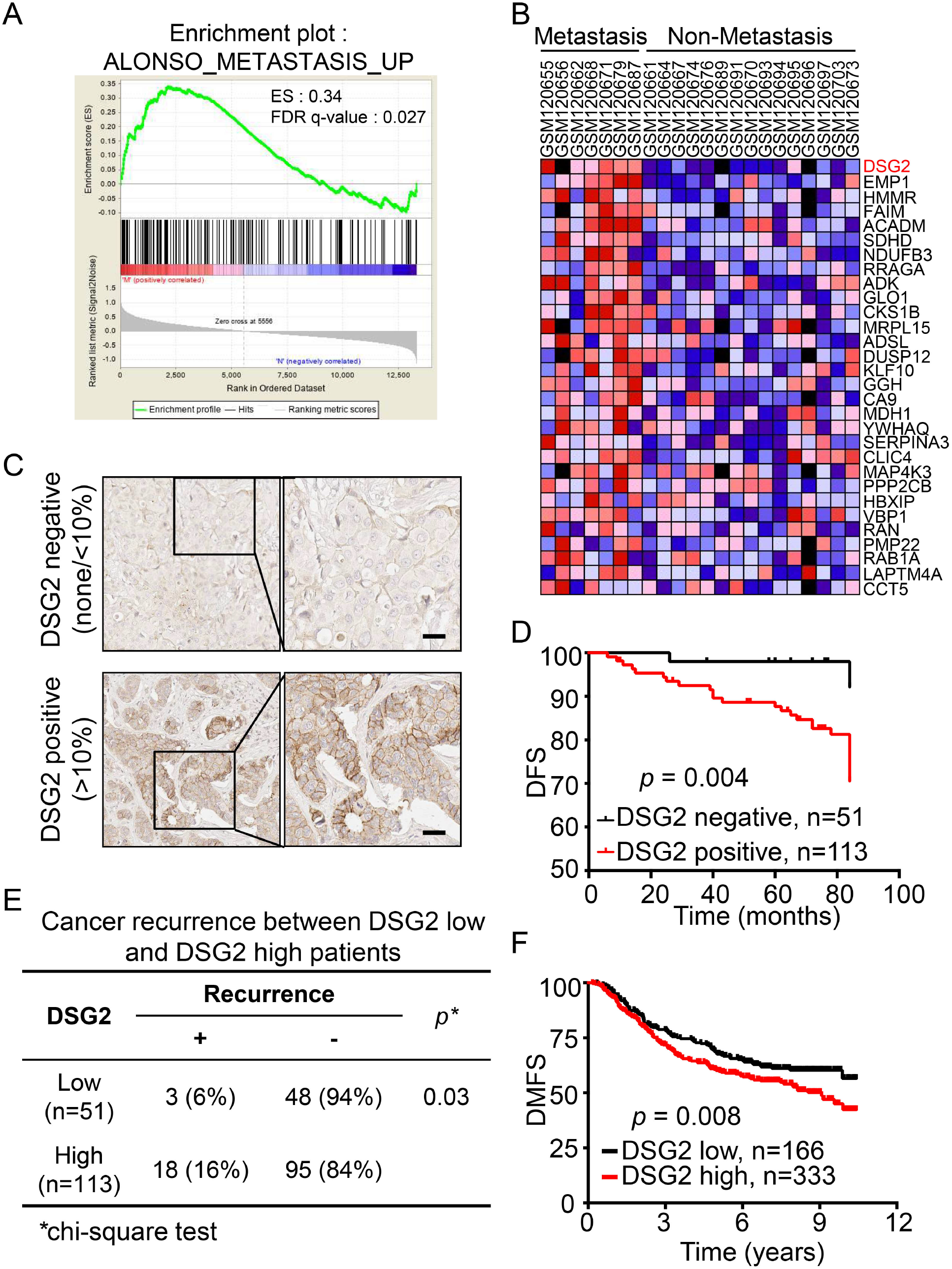
High DSG2 expression is positively correlated with metastasis and high recurrence risk in breast cancer patients. (**A, B**) GSEA-P enrichment plot (A) and heat map (B) for the top 30 up-regulated genes for metastasis in metastatic (n = 7) versus non-metastatic breast cancer patients (n = 15). Normalized enrichment score (NES) and FDR q are listed on the enrichment plots. (**C**) Representative immunohistochemistry (IHC) of breast tumors classified as negative (upper panel; <10% of the tumor cells have detectable DSG2 staining, upper panel) or positive (lower panel; > 10% of tumor cells with intense membrane staining) for DSG2 expression. Enlarged images are presented on the right. Scale bar, 30 μm. (**D**) Kaplan-Meier disease free survival analysis of breast cancer patients grouped by DSG2 expression. DSG2-positive group is indicated by red line (n = 113); DSG2-negative group is indicated by black line (n = 51). *p* = 0.004. The *p*-value was determined by log-rank test. (**E**) Comparison of recurrence rate between patients with DSG2-positive (>10% cells with DSG2 staining) vs. DSG2-negative (<10% with DSG2 staining) tumors using chi-square test. *p* = 0.03. (**F**) Kaplan–Meier analysis of distant metastasis-free survival of patients with different levels of DSG2 using a breast cancer cohort (Yau 2010 dataset) from the UCSC Xena public hub. Receiver operating characteristic (ROC) curve analysis was used to determine the relative level of DSG2. *p* =0.008. The *p*-value was determined by log-rank test.

We then evaluated clinical correlation of DSG2 expression with prognosis using immunohistochemistry (IHC) analysis of 164 biopsy specimens (Fig. 1C). Based on the IHC analysis, patients were grouped into “DSG2-positive” and “DSG2-negative” groups for Kaplan– Meier survival analyses. The disease-free survival was significantly lower in the DSG2-positive group (Fig. 1D, *p* = 0.004). Importantly, DSG2 expression was highly correlated with cancer recurrence (Fig. 1E, *p* = 0.03). To examine whether high DSG2 expression promoted metastasis, we analyzed a breast cancer cohort from the UCSC Xena public hub (Yau 2010 dataset)(24) and found significantly lower distant metastasis-free survival in DSG2-high breast cancer patients (Fig. 1F, *p* = 0.008). The results from these three independent cohorts indicated that DSG2 possesses a metastatic role in breast cancer and could be used as a prognostic marker.

### DSG2 expression is required for CTC clustering and metastasis

To determine DSG2 function in breast cancer progression, we first examined endogenous DSG2 levels in a panel of cell lines (*SI Appendix*, Fig. S2A). MB231, a model cell line for *in vivo* metastasis studies due to its high metastatic potential (25), had the highest level of DSG2 out of eight cell lines assayed. Suppression of DSG2 expression in MB231 using a lentiviral-shRNA system (*SI Appendix*, Fig. S2B) led to depletion of DSG2 from the cell membrane, as confirmed by fluorescence activated cell sorting (FACS) (*SI Appendix*, Fig. S2C). Also, we confirmed that DSG2 was the only highly expressed DSG protein in MB231 and SKBR3 cells (*SI Appendix*, Fig. S2D), and identified shRNAs that specifically depleted DSG2 while not affecting DSG1 or DSG3 (*SI Appendix*, Fig. S2E).

These DSG2-depleted cells were then used to determine the role of DSG2 in CTC clustering and metastasis by orthotopic xenograft assays where EGFP expressing MB231 cells, without (shCtrl) or with (shDSG2) DSG2 depletion, were injected into the 4^th^ mammary fat-pad of NOD/SCIDγmice. Nine weeks after mammary fat pad injection, tumor size, CTC counts and lung nodules were evaluated (Fig. 2A). Mice bearing shCtrl cell-derived, DSG2-expressing, tumors (*SI Appendix*, Fig. S2F) had more metastatic lung nodules compared to the mice bearing shDSG2 cell-derived, DSG2-depleted, tumors (Fig. 2B). Moreover, IHC staining confirmed that the cells from the shCtrl-derived lung tumors expressed DSG2 (Fig. 2B, left panel, *SI Appendix*, Fig. S2F). Interestingly, we also observed a 5-fold higher prevalence of CTC clusters in the shCtrl group (Fig. 2C, 2D, black bars) than the shDSG2 group (278 clusters per gram tumor in the shCtrl group versus 59 clusters per gram tumor in the shDSG2 group). No significant difference in single CTC numbers were found between shCtrl and shDSG2 group (Fig. 2C, 2D, white bars).

**Figure 2.**
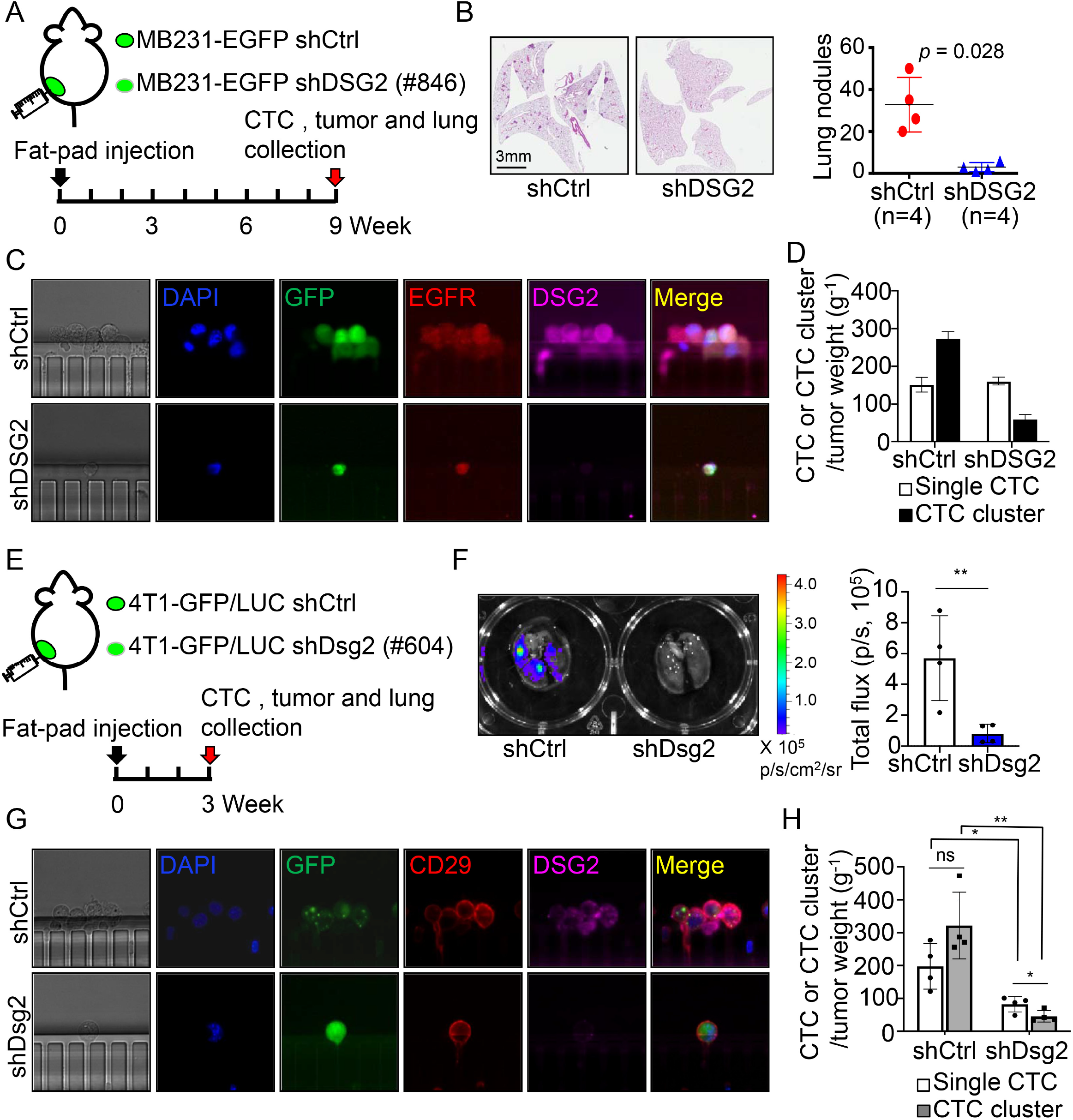
DSG2 expression promotes CTC clustering and metastatic colonization. (**A**) Diagram of the procedure used for orthotopic xenografts. MB231 cells expressing EGFP without (MB231-EGFP shCtrl) or with DSG2 depletion (shDSG2, #846) were injected into 4th mammary fat pads of NOD/SCIDγ mice. CTCs, primary tumors and lung tissues were collected at 9 weeks after injection. (**B**) Representative hematoxylin and eosin (H&E) staining of lung sections from mice orthotopically injected with shCtrl or shDSG2 MB231-EGFP cells and quantification of lung nodule numbers in shCtrl and shDSG2 groups. Error bars indicate S.D., *p* value was determined by unpaired T-test. (**C**) Representative images of CTCs from shCtrl or shDSG2 MB231-EGFP tumor bearing mice captured by MiCareo on-chip filtration and having positive immunofluorescence (IF) staining of DAPI, EGFR and GFP to identify MB231-EGFP cells. (**D**) Numbers of single CTCs and CTC clusters collected from pooled blood of shCtrl or shDSG2 MB231-EGFP tumor bearing mice. Three mice per group were assayed. Data are CTC counts ± 95% confidence intervals. (**E**) Diagram of the syngeneic mouse model procedure. 4Tl-GFP/LUC without (shCtrl) or with DSG2 depletion (shDSG2, #604) were injected into 4th mammary fat pads of BALB/c mice. Primary tumors and lung tissues were collected at 3 weeks after injection. (**F**) Bioluminescence imaging of lung metastasis and quantitation of bioluminescence between shCtrl and shDSG2 groups. Four mice were used for each group. Data are means ± S.D., significant difference determined by T-test. (**G**) Representative images of CTCs from shCtrl or shDSG2 4T1-GFP/LUC tumor bearing mice captured by MiCareo on-chip filtration and having positive IF staining of DAPI, CD29 and GFP to identify 4T1-GFP/LUC cells. (**H**) Numbers of single CTCs and CTC clusters collected from shCtrl- or shDSG2-transduced 4T1-GFP/LUC tumor bearing mice. Four mice were used for each group. Data are means ± SD with significant differences based on unpaired T-test (* indicates *p* < 0.05, ** indicates *p* < 0.01).

A syngeneic tumor mouse model was used to confirm that similar DSG2-driven tumorigenic phenotypes also occurred in immune-competent mice. Mouse mammary tumor cells labeled with GFP-luciferase (4T1-GFP/LUC) with or without DSG2 depletion (*SI Appendix*, Fig. S2G) were injected into the 4^th^ mammary fat-pad of BALB/c mice (Fig. 2E). Consistent with the MB231 xenograft model, shCtrl-injected syngeneic mice had more lung metastasis with DSG2 expressing nodules than shDSG2-injected mice (Fig. 2F, *SI Appendix*, Fig. S2H). Importantly, blood of shCtrl-injected mice also had significantly higher levels of both single CTCs and CTC clusters than blood from the shDSG2 group (Fig. 2G, 2H). This occurred despite the fact that there was no significant difference in primary tumor size between these two groups (*SI Appendix*, Fig. S2I). Together, these data indicated that DSG2 facilitates breast cancer metastasis.

### DSG2 expression promotes metastatic colonization and tumor growth

As colonization is a crucial step to determine whether cancer cells can survive in distant organs (26, 27), we injected SKBR3 cells, which also had high levels of endogenous DSG2 expression (*SI Appendix*, Fig. S2A), with or without DSG2 depletion into mouse tail vein to examine whether DSG2 contributes to colonization ability. Fixation and hematoxylin and eosin (H&E) staining of lung tissue two months after the initial tail vein injection showed that mice injected with DSG2-depleted SKBR3 cells had significantly lower numbers of lung nodules compared to mice injected with control cells (Fig. 3A). Conversely, DSG2 overexpression in MB157 cells, which had low levels of endogenous DSG2 expression (*SI Appendix*, Fig. S2A), significantly increased lung colonization (Fig. 3B).

**Figure 3.**
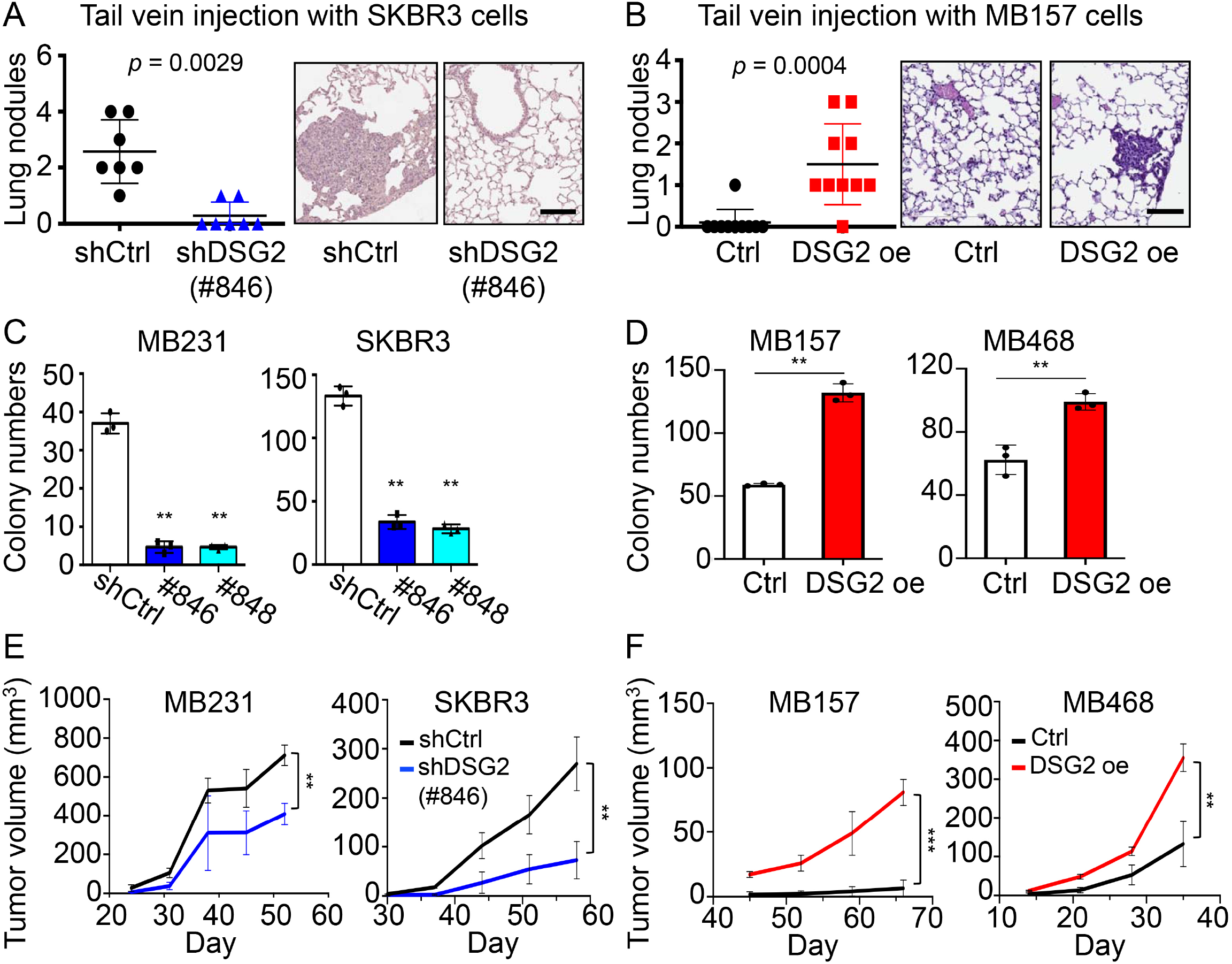
DSG2 expression promoted metastatic colonization and tumor growth. (**A**) Lung metastasis and lung nodule numbers after tail vein injection using control (shCtrl) or DSG2-depleted (#846) SKBR3 cells. Data are means ± S.D. with *p* value based on unpaired T-test. Images show representative H&E staining of lung sections from mice after tail vein injection. Scale bar indicates 100 μm. (**B**) Lung metastasis and lung nodule numbers after tail vein injection using control or DSG2 overexpressing MB157 cells. Data formatting is as described for A. (**C**) Soft agar colony formation assays using DSG2-depleted (#846 or #848) MB231 and SKBR3 cells. Data are shown as mean ± SD with *p* value based on unpaired T-test. The experiments were repeated three times. (**D**) Soft agar colony formation assays using DSG2-overexpressing MB157 and MB468 cells. Replication and data formatting are as described for C. \*\**p* < 0.01. (**E**) Xenograft models in NOD/SCIDγ mice using control (shCtrl) or DSG2-depleted (#846) MB231 and SKBR3 cells. Five mice were used for each group. Data are means ±S.D., significant difference is based on unpaired T-test of the tumor size at the last time point. (**F**) Xenograph models in NOD/SCIDγ mice using control or DSG2-overexpressing MB157 and MB468 cells. Replication and data analysis are as described for E.

In addition, DSG2 promoted tumor growth at the primary site. DSG2 depletion led to significant reduction of colony forming efficiency in soft agar colony forming (SACF) assays (Fig. 3C). Conversely, DSG2 overexpression in MB157 and MB468 cells, which had low endogenous DSG2 levels (*SI Appendix*, Fig. S2A-2C), resulted in higher colony numbers (Fig. 3D). Consistent with the tumorigenic role of DSG2 in promoting colony formation, orthotopic xenografts showed that tumors derived from DSG2-depleted cancer cells were significantly smaller than the control while cells overexpressing DSG2 formed significantly larger tumors (Fig. 3E, 3F, and *SI Appendix*, Fig. S2J, S2K). Together, these results demonstrated that DSG2 expression promotes breast tumor growth at the primary site and increases colonization in the distal organs.

### Repression of DSG2 promotes cell invasion, migration and expression of EMT genes

Initiation of cancer metastasis, including invasion, migration and intravasation, relies heavily on tumor cell dissemination (1, 28). Since DSG2 expression increased CTC clusters and tumor growth in the mammary fat pad and also promoted colonization in the lung, we next evaluated its role in dissemination. Seemingly in contrast to its tumorigenic role, DSG2 knockdown in MB231 cells significantly increased migration ability (*SI Appendix*, Fig. S3A) and DSG2 overexpression in MB157 cells inhibited migration (*SI Appendix*, Fig. S3B). Similarly, depletion of DSG2 increased cell invasion; whereas overexpression of DSG2 inhibited this ability (*SI Appendix*, Fig. S3C, S3D). Consistent with these observations, DSG2 depletion resulted in an upregulation of EMT genes, including *SNAIL*, *SLUG* and *VIMENTIN (VIM)*, all of which are critical for cancer cell mobility and dissemination (29, 30). Conversely, DSG2 overexpression downregulated these genes (*SI Appendix*, Fig. S3E, S3F). Thus, DSG2 suppression may facilitate invasion and migration not only by disrupting cell-cell adhesion but also by upregulating EMT genes to allow cancer cells to detach from the primary tumor and intravasate into the circulatory system as single cells.

### Hypoxia-induced HIF1α suppresses DSG2 expression

Since both single CTCs and CTC clusters were detected in mice with DSG2-expressing tumors (Fig. 2), DSG2 must be suppressed in some parts of the primary tumors to allow cancer cells to invade and disseminate into the blood. To examine this hypothesis, we performed IHC using breast cancer patient samples. A negative correlation of expression pattern between DSG2 and CA9, a hypoxic stress marker, across different tumor regions was observed (Fig. 4; *SI Appendix*, Fig. S4). DSG2 expression was low in hypoxic regions with high CAIX expression (Fig. 4, red box), but high in regions without CAIX staining (Fig. 4, blue box). These results indicated that DSG2 expression may be downregulated under hypoxia.

**Figure 4.**
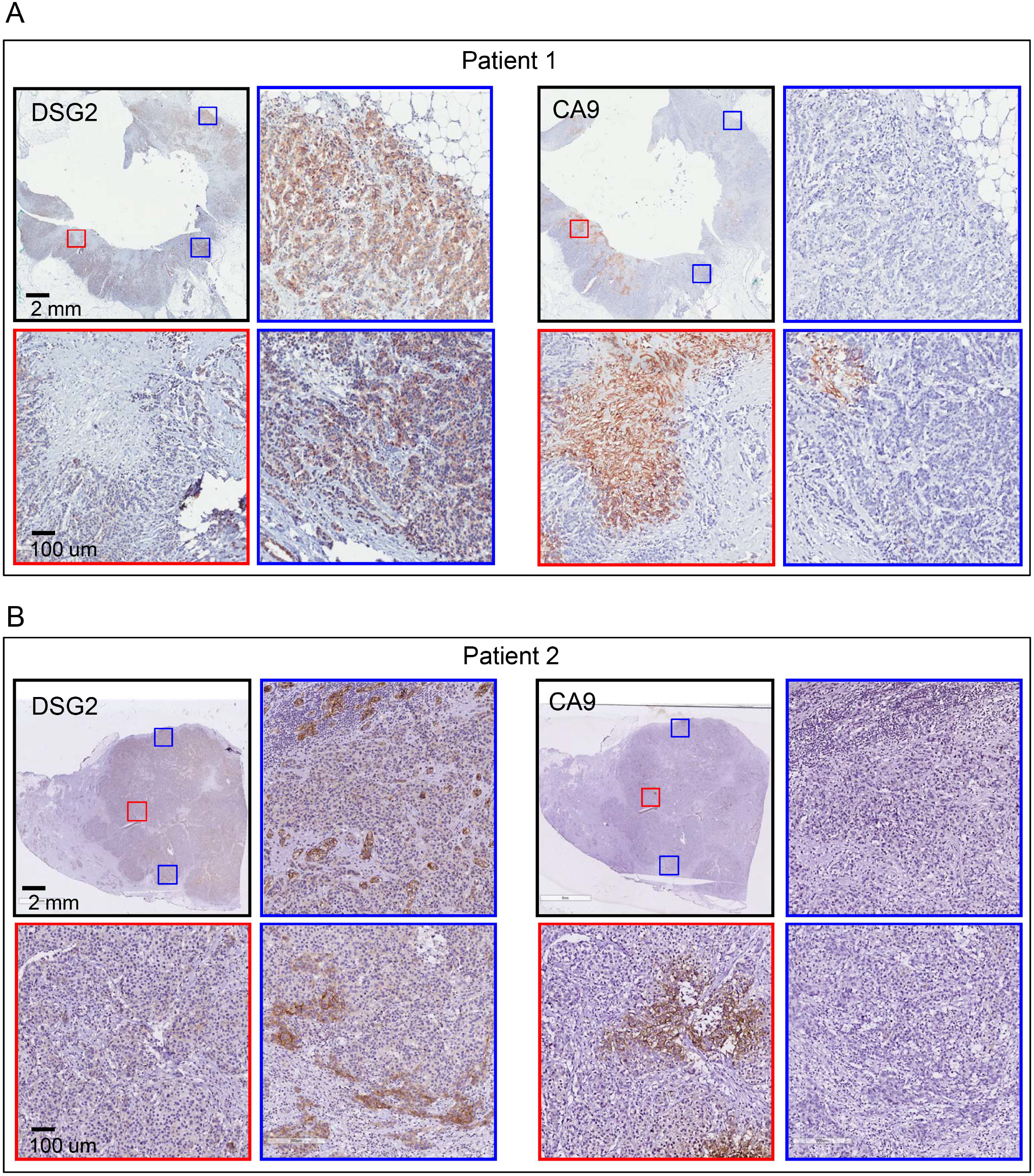
Negative correlation between hypoxia and DSG2 expression observed in clinical specimens. Representative images of DSG2 and CA9 IHC staining using serial tumor sections from clinical breast cancer patients. Red boxes show enlarged images of low DSG2 but high CA9 expression regions. Blue boxes show enlarged images of high DSG2 but low CA9 expression regions. Scale bars indicate 2 mm or 100 μm as shown in the images. Examples shown are representative of 151 patient samples examined. Among the 151 samples, 66 slides contained DSG2^high^/CA9^low^ cancer cells only, 40 slides contained DSG2^low^/CA9^low^ cancer cells only and 45 slides contains both DSG2^high^/CA9^low^ and DSG2^low^/CA9^high^ cells in different regions.

Hypoxia plays a critical role in metastasis (2) but it is not known whether hypoxia affects DSG2 expression. Consistent with the pattern of DSG2 expression in tumor specimens, breast cancer cells subjected to hypoxic stress had decreased DSG2 gene expression and protein level (Fig. 5A, 5B). Similarly, DSG2 expression decreased in cells treated with different doses or durations of the hypoxia-mimetic agent CoCl_2_ (Fig. 5C; *SI Appendix*, Fig. S5A). Since HIF1α is a critical mediator of hypoxia-associated cancer progression (31), we determined the effect of HIF1α on DSG2 expression. Transient transfection of a non-degradable HIF1α (P564A) in 293T cells resulted in downregulation of DSG2 (Fig. 5D). Importantly, depletion of HIF1α in 293T treated with CoCl_2_ abolished hypoxia-induced DSG2 suppression (Fig. 5E). The effect of HIF2α on DSG2 expression was also examined since HIF2α can also be upregulated and contribute to cancer development under hypoxia (32). Unlike HIF1α, expression of non-degradable HIF2α (P531A) did not affect DSG2 expression (*SI Appendix*, Fig. S5B). Moreover, a depletion of HIF2α in SKBR3 did not affect suppression of DSG2 during hypoxia (*SI Appendix*, Fig. S5C). These results indicated that HIF1α is responsible for DSG2 suppression under hypoxic stress.

**Figure 5.**
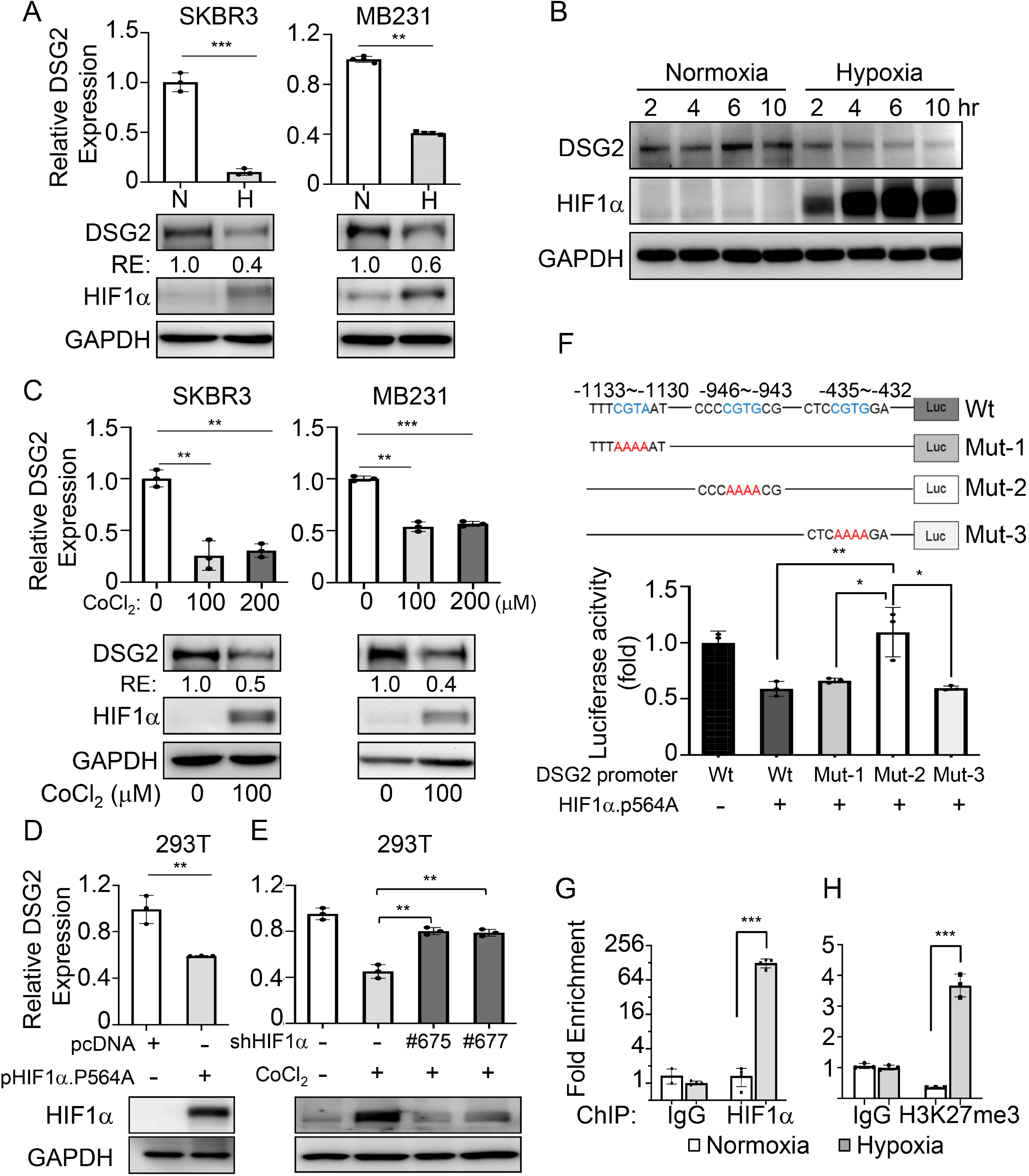
HIF1α suppressed DSG2 expression under hypoxic stress. (**A**) qRT-PCR analysis of *DSG2* level in SKBR3 and MB231 cells under normoxia (N) or hypoxia (H) for 16 hours. Three independent experiments were performed and data are means ± SD from one representative experiment (n =3). Significant differences are based on unpaired T-test. DSG2 and HIF1α protein expression were detected by immunoblot with GAPDH as a loading control. Blots shown are from one representative experiment of three replicates. RE: relative expression level. (**B**) Time-course immunoblot of DSG2 and HIF1α in SKBR3 cells under normoxia or hypoxia. GAPDH was used as a loading control. Blots shown are from one representative experiment of three replicates. (**C**) qRT-PCR analysis of *DSG2* level in SKBR3 and MB231 cells treated with 100 or 200 μM CoCl_2_ for 16 hours. Three independent experiments were performed and data are means ± SD from one representative experiment (n =3). Significant differences are based on unpaired T-test. Elevated HIF1α protein level confirmed that cells were experiencing hypoxic stress. DSG2 and HIF1α expression in cells in the unstressed control or treated with 100 μM CoCl_2_ were detected by immunoblot with GAPDH as a loading control. Blots shown are from one representative experiment of three replicates. RE: relative expression level. (**D**) qRT-PCR analysis of *DSG2* level in 293T cells transiently transfected with pcDNA vector or pcDNA-HIF1α-P564A plasmid. HIF1α expression was detected by immunoblot with GAPDH as a loading control. Three independent experiments were performed and data are means ± SD from one representative experiment (n =3). Significant differences are based on unpaired T-test. (**E**) qRT-PCR of *DSG2* level in 293T cells transduced with shHIF1α lentiviral vectors (clone # 675 or # 677) and treated with 100 μM CoCl2 for 24 hours. Immunoblot of HIF1α shows that hypoxia induced accumulation of HIF1α was effectively blocked by shHIF1α. The experiments were repeated three times. (**F**) Diagram shows three putative HIF1α binding sites on the DSG2 promoter predicted using ISMAR and the mutated promoter sequences used in luciferase reporter assays. Luciferase reporter assays were conducted using 293T cells co-transfected with pcDNA.HIF1α and the wild type (Wt) or mutant DSG2 promoter constructs. Three replicate experiments were performed and data are means ± SD from one representative experiment and significant differences detected using T-test. (**G**) ChIP-qPCR analysis of HIF1α occupancy on the DSG2 promoter region containing putative HIF1α binding site (−946~−943 nt) in SKBR3 cells under normoxia or hypoxia. (**H**) ChIP-qPCR analysis of H3K27me3 on the same DSG2 promoter region as in G. For G and H, three independent experiments were performed and data are means ± SD from one representative experiment with significant differences detected by un-paired T-test. (* indicates *p* < 0.05, ** indicates *p* < 0.01 and *** indicates *p* < 0.001).

To further elucidate whether HIF1α directly suppressed DSG2 transcription, we used Integrated Motif Activity Response Analysis (ISMAR) (33) to identify putative HIF1α binding sites on the DSG2 promoter and found that mutation of one of these sites (−946 to −943) abolished HIF1α inhibition of DSG2 expression (Fig. 5F). Chromatin-immunoprecipitation (ChIP) further confirmed that hypoxia-induced HIF1α bound to this region (Fig. 5G; *SI Appendix*, Fig. S5D) and this HIF1α binding was associated with increased H3K27-trimethylation (H3K27me3), indicative of heterochromatin formation, during hypoxia (Fig. 5H).

This observation of HIF1α-mediated repression of DSG2 contrasted with the more typical function of HIF1α to promote expression of hypoxia-inducible genes (32). To understand how HIF1α carried out this transcriptional suppression, we performed Co-IP assays and found that stabilized HIF1α interacted with polycomb repressive complex 2 components, EZH2 and SUZ12, but not HDAC1 and HDAC2, in breast cancer cells under hypoxia (Fig. 6A, *SI Appendix*, Fig. S5E). This interaction was also confirmed using 293T cells ectopically expressing the non-degradable HIF1α (P564A) (Fig. 6B). Importantly, depletion of EZH2 or SUZ12 significantly abolished hypoxia-induced DSG2 suppression (Fig. 6C) indicating that these PRC2 components are required for DSG2 downregulation under hypoxic stress. Together, these results suggested that HIF1α may recruit EZH2 and SUZ12 to the DSG2 promoter to downregulate DSG2 expression under hypoxia.

**Figure 6.**
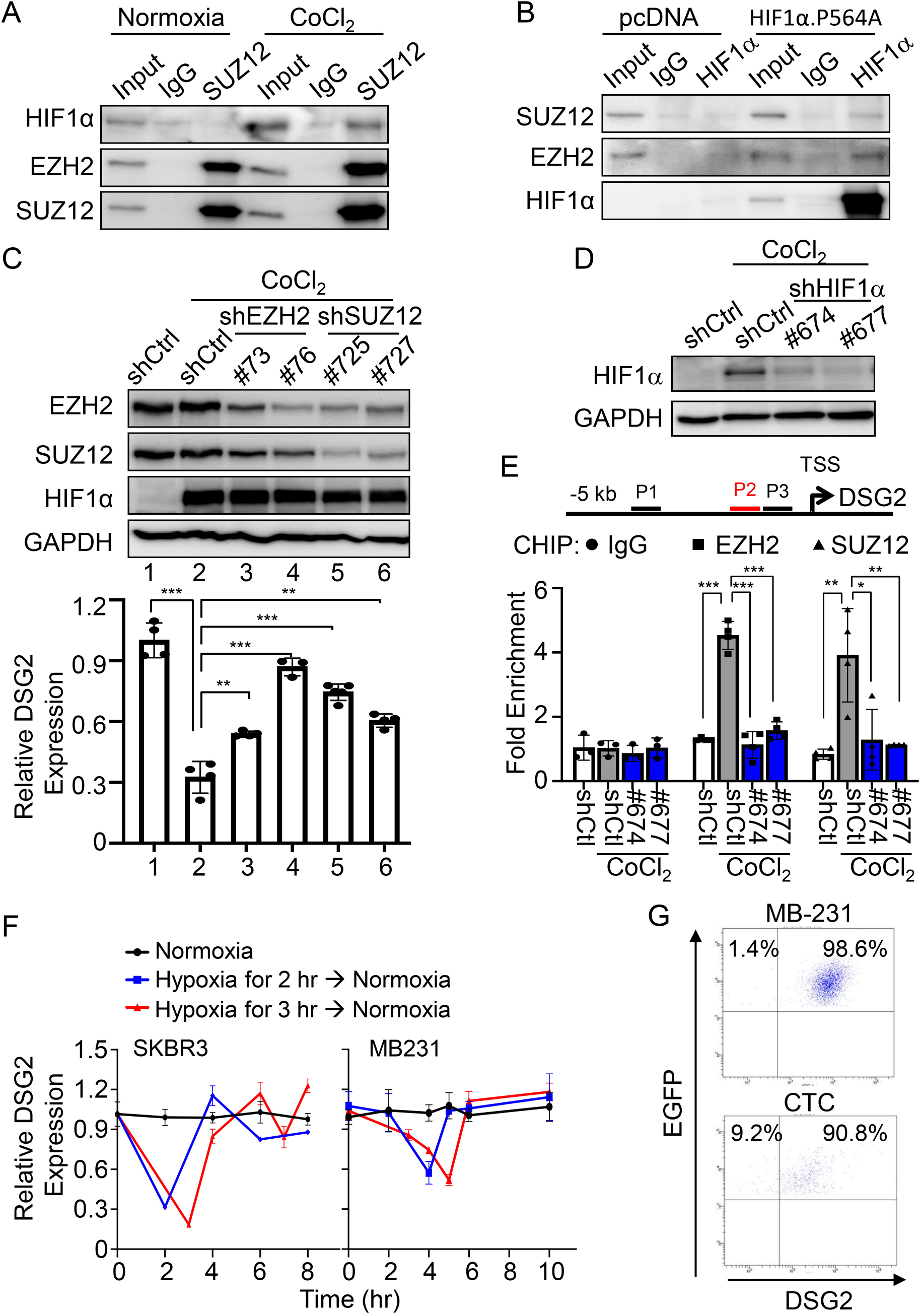
Hypoxia suppresses DSG2 expression via formation of a HIF1α-PRC2 complex. (**A**) Co-IP of HIF1α, EZH2 and SUZ12 in SKBR3 cells without (normoxia) or with CoCl_2_ treatment. IgG was used as a negative control. The experiments were repeated three times with consistent results. (**B**) Co-IP of SUZ12, EZH2 and HIF1α in 293T cells transfected with HIF1α.P564A. IgG was used as a negative control. The experiments were repeated three times. (**C**) Immunoblot of EZH2, SUZ12 and HIF1α (top) and qRT-PCR analysis of DSG2 level (bottom) in shCtrl (shC), shEZH2 or shSUZ12-transduced SKBR3 cells treated with CoCl_2_. GAPDH was used as a control. qRT-PCR data are means ±S.D. (n = 3). Significant differences were detected by T-test. (**D**) Immunoblot of HIF1α in SKBR3 cells transduced with shHIF1α lentiviral vectors (clone # 674 and # 677) and treated with CoCl_2_ for 6 hours. The experiments were repeated three times. (**E**) ChIP-qPCR analysis of EZH2 and SUZ12 on the P2 promoter region (−1062 to −552) in HIF1α-depleted SKBR3 cells with or without CoCl_2_ treatment. Data are mean ± SD with significant differences base on unpaired T-test (\**p* < 0.05, \*\**p* < 0.01. \*\*\**p* < 0.001). The experiments were repeated three times. (**F**) qRT-PCR analysis of DSG2 in SKBR3 (left panel) and MB231 (right panel) cells kept under normoxia throughout the experiment (black line), or under hypoxia for 2 hr (blue line) or 3 hr (red line) and then released from the stress. Data are means ±S.D (n = 3). Significant differences are based on T-test of data at each time point. The experiments were repeated three times. (**G**) FACS analysis of DSG2 levels in shCtrl EGFP-MB231 cells before injection (top panel) or in CTCs collected from pooled blood of four mice at 9 weeks after injection (bottom panel).

To further investigate whether EZH2 and SUZ12 were recruited to the DSG2 promoter by HIF1α under hypoxic stress, we performed ChIP assays using cells with or without HIF1α depletion (Fig. 6D). Upon CoCl_2_ treatment, HIF1α was induced and recruited these co-repressors to the HIF1α-binding region on the DSG2 promoter in the shCtrl cells (Fig. 6E, gray bar). However, the recruitment of EZH2 or SUZ12 was abolished in HIF1α depleted cells under hypoxic stress (Fig. 6E, blue bar). These results demonstrated that stabilized HIF1α is required to recruit PRC2 complex to the DSG2 promoter to suppress DSG2 transcription under hypoxia.

Since DSG2 is critical for distant organ colonization and is highly expressed in metastatic nodules (Fig. 2 and Fig. 3), DSG2 suppression that occurs in hypoxic conditions must be released before colonization can occur. Consistent with this idea, de-repression of DSG2 expression was observed within one hour after SKBR3 cells were released from hypoxia *in vitro* (Fig. 6F, left panel). Interestingly, the expression of DSG2 continued decreasing for approximately one hour after MB231 cells were released from hypoxic stress (Fig. 6F, right panel). Nevertheless, reactivation of DSG2 expression occurred soon after that. To confirm that this reactivation is important *in vivo*, we injected EGFP expressing MB231 cells into mammary fat pads and evaluated DSG2 expression in CTCs after injection. Before injection, nearly all cells expressed DSG2 (98.6%, Fig. 6G, top panel). At nine weeks after injection, more than 90% of CTCs disseminated from the primary tumor site expressed DSG2 while 9% had low/non-detectable DSG2. Given our other observations, it seems likely that the cells with low or non-detectable DSG2 had just been released from the primary tumor and undergone intravasation (Fig. 6G, bottom panel). Taken together, our *in vitro* and *in vivo* observations are all consistent with dynamic regulation of DSG2 where hypoxia-mediated DSG2 suppression followed by de-repression of DSG2 when hypoxic stress is released in the circulatory system is required for DSG2-promoted breast cancer metastasis.

## Discussion

Our study demonstrated that dynamic changes in DSG2 expression are important for breast tumorigenesis and malignancy (Fig. 7). Expression of DSG2 not only facilitated breast tumor growth in mammary tissue and colonization in lung, it also promoted CTC collective migration in the circulatory system (Fig. 2, 3). In hypoxic tumor regions, HIF1α-mediated DSG2 suppression was critical for cancer cells to gain mobility, detach from the primary tumor and intravasate into the blood (Fig. 7; *SI Appendix*, Fig. S3).

**Figure 7.**
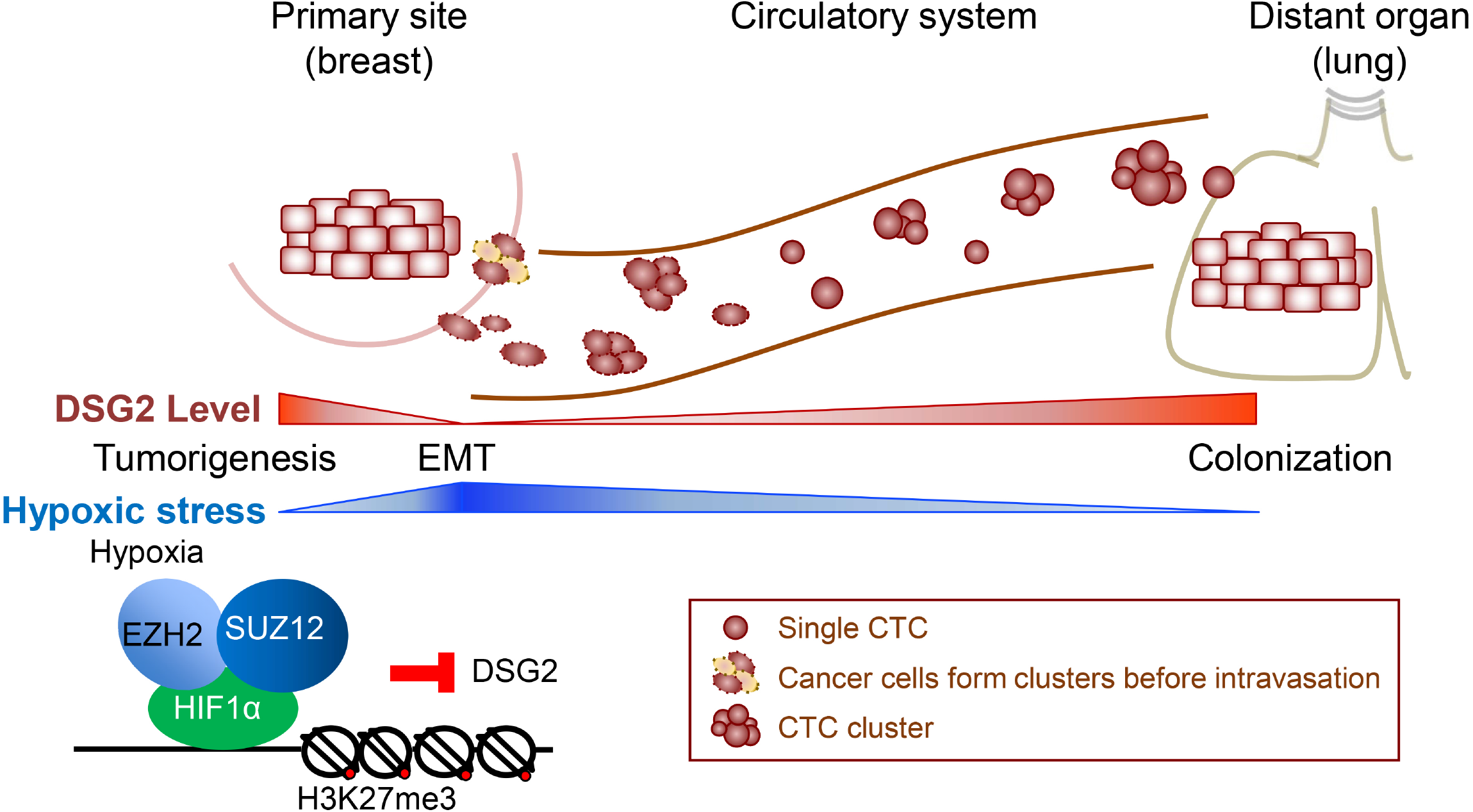
Dynamic changes of DSG2 expression contribute to breast tumorigenesis and metastasis. High DSG2 expression promotes breast tumor growth in mammary tissue. In non-hypoxic tumor regions with high DSG2, CTC clusters may be shed and DSG2-mediated cell adhesion may facilitate CTC clustering prior to intravasation. Alternatively, when the tumor is large enough to induce hypoxia, HIF1α is stabilized and recruits PRC2 complex (EZH2 and SUZ12) to the DSG2 promoter to suppress its transcription. DSG2 suppression allows cancer cells to undergo EMT and single CTCs are released from the primary tumor. In circulatory system, hypoxic stress is released and reactivation of DSG2 expression allows colonization.

Recent research has proposed two mechanisms describing how CTCs form clusters to collectively migrate in the circulatory system. One is that tumor cells intravasate as multicellular clusters via plakoglobin-dependent intercellular adhesion into the blood (9). The other is that individual tumor cells aggregate via CD44-PAK2 mediated FAK signaling after intravasation (15). Since DSG2 is an integral component of desmosomes and its cytoplasmic tail binds to plakoglobin and plakophilins (34), DSG2 may facilitate CTC clustering prior to intravasation through its cell-cell adhesion function. Therefore, non-hypoxic tumor regions where DSG2 expression is high may shed CTC clusters.

In contrast, hypoxic tumor regions where DSG2 is downregulated may more readily generate single CTCs. CTCs that can survive in the circulatory system and colonize in distant organs are thought to be cancer stem-like cells and are highly associated with metastasis (35, 36). We also observed that DSG2 expression promoted mammosphere formation and survival in a low-attached culture system (*SI Appendix*, Fig. S6), suggesting that DSG2 may contribute to CTC survival during circulation and may help maintain cancer stemness. Consistent with this notion, DSG2 has recently been identified as a marker for pluripotent stem cells (PSCs) and is critical for PSC self-renewal and reprogramming (37). These observations could partially explain why there was no significant difference or even higher single CTC numbers in mice bearing cancer cells with high overall DSG2 level (shCtrl) compared to mice bearing DSG2-depleted (shDSG2) cancer cells (Fig. 2D, 2H). The shCtrl group may have increased single CTCs due to intratumoral hypoxia which locally repressed DSG2 expression. Since these shCtrl-single CTCs are able to reactivate DSG2 expression when released from hypoxia, they may survive better in the blood. However, the single CTCs generated from the shDSG2 tumors cannot reactivate DSG2 expression and thus may have reduced survival. Taken together, high DSG2 expression may allow CTC clusters to remain intact and single CTCs to persist longer in the circulatory system. Both of these scenarios can increase the chance for colonization. It is also possible that DSG2 directly promotes colonization and metastatic tumor growth via its roles in cell adhesion.

In several cancers, HIF1α acts as a transcriptional activator to upregulate genes required for survival, EMT, angiogenesis and metastasis upon hypoxic stress (31, 32). HIF1α upregulates these target genes by recruiting coactivators such as p300/CBP acetyl-transferase, CDK8-mediator, Potin and SWI/SNF chromatin remodelers (38). In contrast, there is relatively little evidence that HIF1α can also act as a transcription repressor. Under hypoxia, stabilized HIF1α was found to displace the transcription activator Myc from Sp1 binding to repress MutSα expression to contribute to genomic instability in colon cancer cells (39). However, what corepressors HIF1α recruited to the target promoter upon hypoxia was not known. Our observation that HIF1α recruited EZH2 and SUZ12 to suppress DSG2 transcription (Fig. 5, 6) provides a new regulatory mechanism and expands the oncogenic role of HIF1α in promoting breast cancer progression. Whether this HIF1α-EZH2-SUZ12 complex serves as a common gene suppression mechanism to facilitate cancer malignancy in other cancer types is worthy of investigation.

Our clinical data and animal models clearly demonstrated that DSG2 can be used as a prognostic marker for breast cancer. Interestingly, DSG2 expression is also associated with poor prognosis in other cancers including cervical, head and neck and lung cancers (*SI Appendix*, Fig. S7). This suggests a widespread role of DSG2 in cancer malignancy. As a transmembrane protein, DSG2 may be a drugable target. However, the strategy has to be carefully designed. Using a monoclonal DSG2 specific antibody to target the DSG2 to inhibit anchorage independent growth may be possible. However, antibody-DSG2 internalization may occur and result in enhancement of intravasation, as downregulation of DSG2 promotes invasion. In this case, a well-designed antibody-cytotoxic drug conjugate (ADC) (40) may be a more effective therapeutic method to target DSG2 expressing cancer cells. In addition, since DSG2 expressing CTCs tend to form clusters which facilitate distant organ colonization, dialysis for DSG2 expressing cancer cells in the blood may be another way to inhibit metastasis. Alternatively, DSG2 may be used as a specific marker for CTC cluster detection and isolation. Both the underlying molecular mechanisms of DSG2-mediated tumorigenesis and the possibility of using DSG2 as a cancer therapeutic target are promising areas for further work.

## Materials and Methods

Breast cancer specimens for IHC analysis were collected from National Taiwan University Hospital. All specimens were encoded to protect patients under protocols approved by the Institutional Review Board of Human Subjects Research Ethics Committee of Academia Sinica (AS-IRB01-16031) and National Taiwan University (201605057RINA), Taipei, Taiwan. Animal care and experiments were approved by the Institutional Animal Care and Utilization Committee of Academia Sinica (IACUC# 15-11-885). NOD/SCID mice were kindly provided by Dr. Michael Hsiao (Genomics Research Center, Academia Sinica, Taiwan) and BALB/c mice were purchased from the National Laboratory Animal Center (Taiwan). Cancer cell lines including MB231, MB157, MB468, SKBR3 and 4T1 were obtained from the American Type Culture Collection. Human breast cancer cell lines were maintained in DMEM and mouse breast cancer cell line 4T1 was maintained in RPMI supplemented with 10% fetal bovine serum and antibiotics and cultured at 37°C in a humidified incubator supplemented with 5% CO_2_. For hypoxia experiments, medium was first incubated in HypoxyCOOL system (Baker) for 8 hours to reduce oxygen level to 1%. This medium was then used to culture the cells in INVIVO 400 hypoxia chamber (Baker) supplemented with 5% CO_2_ and 1% O_2_ at 37°C. Single CTC and clusters were identified using a CTC platform from MiCareo, Inc. A spike-in control experiment using 10^3^ 4T1-GFP/LUC cells mixed with 2ml blood from wildtype BALB/c mice was performed to determine whether the centrifugation step of CTC isolation increases cell aggregation (*SI Appendix,* Fig S8). Further details of experimental methods are given in the *SI Appendix*, Materials and Methods.

## Author contributions

P.-H.C. and W.W.H.-V. conceived the study. P.-H.C. performed all experiments not attributed to other authors. P.-H.H. assisted with the LC-MS/MS analysis in the original manuscript and Y.-M.J. assisted with immunohistochemistry analysis, P.-H.C., G.Y.T.T. and P.-H.H. analyzed the data, P.-H.C., M.-C.C. and Y.-P.T. generated the DSG2 antibody, Y.-M.J., Y.-F.T. and M.-H.Y. provided clinical samples. P.-H.C. and W.W.H.-V. wrote the paper.

## Acknowledgements

This work was supported by Academia Sinica [AS-SUMMIT-108, AS-SUMMIT-109] and the Taiwan Ministry of Science and Technology [MOST 109-0210-01-18-02, MOST 108-3114-Y-001-002, MOST 107-0210-01-19-01, MOST 105-2628-B-001-008-MY3 and MOST-108-2311-B-001-005-MY3]. The authors would like to thank Dr. Michael Hsiao (Genomics Research Center, Academia Sinica, Taiwan) for providing NOD/SCID mice, and Dr Yi-Cheng Chang (Graduate Institute of Medical Genomics and Proteomics, National Taiwan University) for discussion.

