## Supplemental Methods, Figures and Tables for "Interplay between Desmoglein2 and hypoxia controls metastasis in breast cancer"

**This PDF file includes:**

Supplementary text: Materials and Methods
Figures S1 to S8
Tables S1 to S2
SI References

### Supplementary text: Materials and Methods

### Materials and Methods

#### Cell lines and cell culture

Cancer cell lines including MB231, MB157, MB468, SKBR3 and 4T1 were obtained from the American Type Culture Collection. Human breast cancer cell lines were maintained in DMEM and mouse breast cancer cell line 4T1 was maintained in RPMI supplemented with 10% fetal bovine serum and antibiotics and cultured at 37°C in a humidified incubator supplemented with 5% $CO_2$. For hypoxia experiments, medium was first incubated in HypoxyCOOL system (Baker) for 8 hours to reduce oxygen level to 1%. This medium was then used to culture the cells in INVIVO 400 hypoxia chamber (Baker) supplemented with 5% $CO_2$ and 1% $O_2$ at 37°C.

#### Gene Set Enrichment Analysis (GSEA)

Transcriptome data of 7 metastatic and 15 non-metastatic breast cancer patients of a breast cancer cohort (GSE5327) from the GEO (http://www.ncbi.nlm.nih.gov/geo/) database (1) was used. GSEA was performed with metastasis (ALONSO_METASTASIS_UP) (2) or cell junction (GO_CELL_CELL_Junction, Gene Ontology) gene sets. The Rank Metric Score of genes listed in Figure 1 were all above 0.25. The enrichment score (ES) of a gene set with a FDR q value below 0.2 was considered to be significantly enriched.

#### Quantitative real-time PCR (qRT-PCR) and immunoblot (IB)

Total RNA was extracted using TRI reagent (Sigma-Aldrich, Cat. T9424) and quantified by NanoDrop (ThermoFisher). cDNA was reverse transcribed with QuantiNova Reverse Transcription Kit (Qiagen, Cat. 205410). qRT-PCR was performed using an ABI Step-

One Plus instrument with SYBR-Green reagents (Applied Biosystems) according to the manufacture's instruction with gene specific primers (Appendix Table 1). The relative quantities of mRNAs were determined using comparative cycle threshold methods, and were normalized against the mRNA of *GAPDH*.

For IB, whole cell lysate was prepared using RIPA lysis buffer (25 mM Tris-HCl pH 7.6, 150 mM NaCl, 5 mM EDTA, 1% NP-40, 1% sodium deoxycholate, and 0.1% SDS) with 1 mM PMSF, protease inhibitor cocktail tablet (Sigma-Aldrich, Cat. S8830), and phosphatase inhibitor cocktail (Roche, Cat. 04906837001), followed by sonication at 4°C using UP50H (Hielscher). Protein concentration was determined by Bradford assay (Genecopoeia, Cat. P010). Immunoblot analysis was performed after SDS-PAGE, with overnight incubation with a 1:1000 dilution of primary antibody against DSG2, HIF1α and GAPDH (Appendix Table 2) and followed by a 1:10000 dilution of horseradish peroxidase-conjugated anti-rabbit or anti-mouse antibody (Bethyl, Cat. A120-102P and Cat. A90-516D2). Signals were detected using Western Chemiluminescent HRP Substrate (Millipore, Cat. WBKLS0500). At least three independent experiments were performed.

**Generation of monoclonal anti-human DSG2 antibody**

Hybridoma methodology was employed for the development of monoclonal antibodies (mAbs) specific to DSG2. The antigen was a recombinant fusion protein containing the DSG2 extracellular domain 2 (EC2, 160-273 a.a) expressed in the E. coli strain BL21. To induce anti-DSG2 immune response, BALB/c mice were immunized twice subcutaneously with 50 μg of DSG2 EC2 recombinant proteins with TiterMax Gold adjuvant (Sigma-Aldrich, Cat. T2684) according to the manufacturer's instruction.

Hybridoma supernatants were used to screen anti-DSG2 mAbs for binding to DSG2 EC2 proteins by enzyme-linked immunosorbent assay (ELISA).

**Immunohistochemistry (IHC)**

Formalin-fixed paraffin embedded primary and xenograft tumor tissue sections were used for IHC. Heat induced antigen retrieval was performed using Trilogy buffer (Cell Marque, Cat. 920P) at 95°C for 20 min. Endogenous peroxidase was eliminated with 3% $H_2O_2$ for 10 mins. Slides were blocked with PBS containing 10% FBS for 1hr at room temperature, followed by primary antibodies against DSG2 and CAIX (Genetex, Cat. 70020 ) in PBS/10% FBS overnight at 4°C. After washing, slides were incubated with HRP rabbit/mouse polymer for 1 hr before visualization with liquid diaminobenzidine tetrahydrochloride plus substrate DAB chromogen from Dako REAL EnVision (Dako, Cat. K5007). All slides were counterstained with hematoxylin. The images were captured by an Aperio Digital Pathology system. IHC sample was scored as “negative” if tissue section containing all cells with no staining or cell membrane staining is observed in <10% of the tumor cells. Sample was scored as “positive” if tissue section containing more than 10% of tumor cells with intense membrane staining.

**Soft agar colony formation assay**

Two thousand cells were mixed with a layer of 0.35% agar/complete growth medium over a layer of 0.5% agar/complete growth medium in a well of a 12-well plate. After culture at 37°C for 2-3 weeks, crystal violet-stained colonies were counted.

**Invasion and migration assay**

For invasion assay(3), ten thousand cells were seeded in a 24-well transwell insert (Falcon HTS Fluoro Block; pore size, 8 μm) coated with Matrigel (Corning, Cat. 354234) in growth medium without serum. DMEM with 10% FBS was used as a chemoattractant in the lower chamber of 24-well plate. After 16-24 hr incubation, the invaded cells were fixed with methanol and stained with DAPI. Cells were counted with fluorescence microscopy.

The scratch assay(4) was performed with $10^5$ cells in a well of a 6-well plate. After the cell density reached 100% confluence, a vertical wound on the cell monolayer was made using a micropipette. The medium and cell debris were removed and the cell monolayer was washed with PBS. Fresh medium was then added and the cells were cultured for additional 12-32 hrs.

**Plasmids and reagents**

The lentiviral pLKO-puro-shRNA expression vectors shCtrl (TRCN0000072224), shDSG2 [TRCN0000053846 (#846) and TRCN0000289848 (#848)], shDsg2 [TRCN0000094604 (#604) and TRCN0000094608 (#608)], shEZH2 [TRCN0000040073 (#73) and TRCN0000040076 (#76)], shSUZ12 [TRCN0000038725 (#725) and TRCN0000038727 (#727)], shHIF1α [TRCN0000318674 (#674), TRCN0000318675 (#675) and TRCN0000318677 (#677)], shHIF2α [TRCN000033342501 (#501) and TRCN0000352630 (#630)] and EGFP plasmid (pAS7w) were purchased from the National RNAi Core Facility (Taipei, Taiwan). The non-degradable pHIF1α.P564A and pHIF2 α.P531A plasmid was kindly provided by Dr. Hsiu-Ming Shih (Academia Sinica). DSG2 promoter luciferase reporter plasmid was purchased from GeneCopoeia (HPRM23502-LvPG04). Site-directed mutagenesis was performed to create mutation in three regions of the DSG2 promoter (-433 to -436, -944 to -947 and -1131 to -1134).

**Lentivirus expression system**

Lentivirus packaging was done by co-transfection of pMD.G, pCMVR8.91 and pLKO-puro-shRNA in 293T cells using LT1 transfection reagent (Mirus Bio, Cat.MIR 2304). *DSG2* was cloned into a lenti-viral pLVX-IRES-Neo bi-cistronic lentiviral expression vector (Clontech, Cat. 631251) using XhoI/NotI (pLVX.DSG2).

**Co-immunoprecipitation (Co-IP)**

For CoIP assays, DSP (Thermo, Cat.22585) was used to crosslink interacting proteins. After crosslink, whole cell lysate was obtained using RIPA buffer supplemented with PMSF, protease inhibitor cocktail (Sigma-Aldrich, Cat.S8830) and phosphostop (Roche, Cat. 4906845001) followed by sonication and centrifugation at 12,000 × g at 4°C for 10 min. Normal rabbit IgG and A/G agarose (Thermo, Cat.20421) were used to pre-clean the cell lysate before immunoprecipitation with antibodies against DSG2, HIF1α, EZH2, SUZ12 or normal rabbit IgG overnight at 4°C with gentle agitation. Then, 40 µl prewashed protein A/G agarose were added to the mixture and incubated at room temperature for 2 h with gentle agitation. After extensive washing with IP buffer (20mM HEPES, 20% glycerol, 2% NP-40, 300mM NaCl, 1% sodium deoxycholate and 0.2% SDS) twice and PBS, interacting proteins were eluted with SDS-PAGE. Immunoblot was performed with anti-HIF1α (Genetex, Cat.127309), SUZ12 (Cell signaling, Cat.3737), EZH2 (Cell signaling, Cat.5246), HDAC1 (Cell signaling, Cat.5356), SP1 (Cell signaling, Cat.5931), EGFR (Cell signaling, Cat.4267), HDAC2 (Cell signaling, Cat.2545) and Tri-methyl-histone H3K27 (Cell signaling, Cat.9756) antibodies (see Appendix Table 2 for detailed information).

**Chromatin immunoprecipitation (CHIP)**

Chromatin immunoprecipitation assay was performed as previously reported (3). Cells were cross-linked with 1% formaldehyde for 10 min at 37°C and the crosslinking was stopped by 1.25 M Glycine. Cross-linked chromatin was sonicated to a size with a range between 150 to 350 bases. Immunoprecipitations were performed with antibodies against EZH2, SUZ12, Tri-methyl-histone H3K27 and HIF1α (Appendix Table 2) and corresponding control (immunoglobulin G) antibodies and Protein A/G Agarose. Semi-quantitative PCR was performed to detect protein associated promoter regions using primers listed in (Appendix Table 1).

**Promoter luciferase assay**

For 293T cells, $1x10^5$ cells were seeded in a well of a 12-well plate and co-transfected with wild type or mutant DSG2, Renilla.Luc (for normalization), and HIF1α.P564A or pcDNA vector using LT1 transfection reagent for 8 hr. Cells were then cultured in fresh medium for 24 hr. Cell extracts were collected and the luciferase activity was measured using Secrete-Pair Dual Luminescence Assay Kit (GeneCopoeia, Cat. LF031) according to the manufacture's instruction.

**Mouse tumorigenic and colonization assay**

Animal care and experiments were approved by the Institutional Animal Care and Utilization Committee of Academia Sinica (IACUC# 15-11-885). Non-obese diabetic/severe combined immunodeficient mice (NOD/SCID) were kindly provided by Dr. Michael Hsiao (Genomics Research Center, Academia Sinica, Taiwan). NOD/SCID fat pads were injected with $2\times10^6$ cancer cells transduced with shCtrl, shDSG2 or pLVX.DSG2 lentiviral vector mixed with Matrigel (1:1). Tumor volumes were evaluated every 7 days after injection. Mice were sacrificed two months after injection and the tumors were weighed and volumes were measured for tumorigenesis evaluation. In the 4T1 syngeneic model, BALB/c mice were injected with 2 x $10^5$ 4T1-GFP/LUC cells transduced with shCtrl or shDsg2 lentiviral vector mixed with Matrigel. Mice were sacrificed 3 weeks after injection. IVIS kinetics imaging system (Caliper LifeSciences) was applied to monitor tumor metastasis. For colonization assay, $10^5$ cancer cells transduced with shCtrl, shDSG2, pLVX.Ctrl or pLVX.DSG2 lentiviral vector were injected intravenously. Two months after injection, mice were sacrificed and lungs were collected. Lungs were fixed and sectioned. Numbers of metastatic lung nodules were identified and counted using H&E staining.

**Flow cytometry (FACS)**

Cancer cell lines including MB231, MB157, MB468, and SKBR3 were trypsined with trypsin buffer (1 x HBSS without $Ca^{2+}$ and $Mg^{2+}$ containing 5 mM EDTA). Cells were washed with PBS and resuspended in staining buffer (2% FBS with 2 mM EDTA in PBS). DSG2 antibody was used to stain cells on ice for 30 mins. After centrifugation at 500×*g* for 5 min, cells were washed twice with PBS and then stained with anti-mouse 488 (Invitrogen, Cat.A28175). After washing with PBS twice, cells were determined by FACS analysis.

**Detection of circulating tumor cells (CTCs)**

Single CTC and clusters were identified using a CTC platform from MiCareo, Inc. NOD/SCID mouse fat pads were injected with 2 x $10^5$ MB-231 cells transduced with shCtrl or shDSG2 (4 mice each group). Five weeks after injection, 2.3 ml blood from each group was collected and transferred into a $K_2$EDTA tube (BD, Cat.367835).Samples were incubated with PE-conjugated-anti-EGFR antibody for 20 minutes at room temperature and then diluted in 27 ml ISOTON Diluent (Beckman, Cat. 8546719). After centrifugation, supernatant was removed and the samples within each group were pooled for CTC analysis. A cell was classified as a CTC by its morphological features, positive staining of DAPI, and expression of EGFR and GFP. Aggregation of cells containing more than three CTCs is defined as a CTC cluster. For syngeneic model, BALB/c mice were injected with 2 $x10^5$ 4T1-GFP/LUC cells. Three weeks after injection, 0.8 ml blood per mouse was collected for CTC analysis. To determine whether the centrifugation step increases cell aggregation, a spike-in control experiment using $10^3$ 4T1-GFP/LUC cells mixed with 2ml blood from wildtype BALB/c mice was performed (Fig S8). To determine DSG2 levels in CTCs disseminated from the EGFP-labeled MB231 shCtrl tumor, ~2 ml blood from 3 mice was collected (0.7 ml blood each mouse). The blood samples were diluted 1:10 in ACK lysing buffer ($NH_4Cl$, $KHCO_3$, $EDTANa_22H_2O$) for 5 min to lyse red blood cells (RBCs). $CD45^+$ immune cells were removed using anti-CD45 antibody (Invitrogen, Cat.14045182) and magnetic beads on a DynaMag™-2 magnet (Invitrogen, Cat. 12321D). The supernatant was collected and the remaining cells were stained with anti-DSG2 antibody followed by anti-mouse 647 secondary antibody (Sigma-Aldrich, Cat.SAB4600182). DSG2 levels were determined in $EGFP^+$ population by FACS analysis. Blood from mice without tumor innoculation was used as a negative control.

**Statistics**

All data were presented in replicates of three or more and presented as means ± SD. Student’s t-test was used to compare control and treatment groups. All statistical analyses were performed using Prism 8 software. For migration assay, the size of wounds was measured using snapshot picture by microscopy. For clinical correlation, the Kaplan-Meier estimation method was used for overall survival analysis, and a log-rank test was used to compare differences. For the data acquired from the UCSC Xena public hub (Yau 2010 dataset), receiver operating characteristic (ROC) curve analysis was used to classify the level of *DSG2*. No violation of the proportional assumption was detected.

**Study approval**

Breast cancer specimens for IHC analysis presented in Fig. 1 and Supplementary Fig. 1 were collected from National Taiwan University Hospital. All specimens were encoded to protect patients under protocols approved by the Institutional Review Board of Human Subjects Research Ethics Committee of Academia Sinica (AS-IRB01-16031) and National Taiwan University (201605057RINA), Taipei, Taiwan.

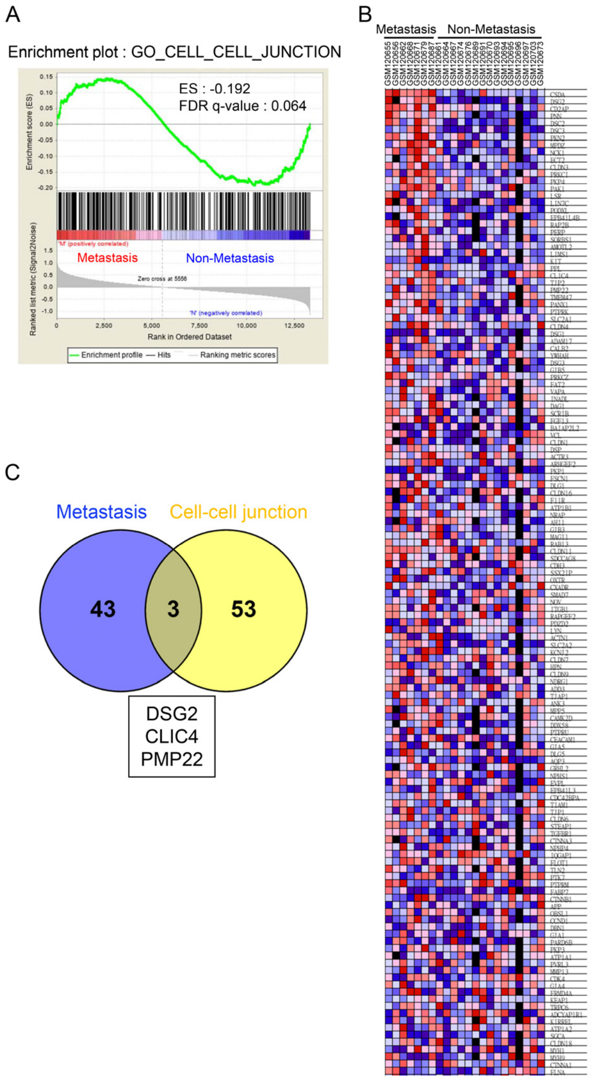


**Figure S1**

**(A, B)** GSEA (A) and heat map (B) for the top 100 up-regulated genes in metastatic (n = 7) versus non-metastatic breast cancer patients (n = 15) using cell-cell junction gene set. Normalized enrichment score (NES) and FDR q are listed on the enrichment plots. **(C)** Venn diagram showing overlap of the metastasis and cell-cell junction gene datasets. (Venn diagrams constructed using the venny online tool (https://bioinfogp.cnb.csic.es/tools/venny/index.html)

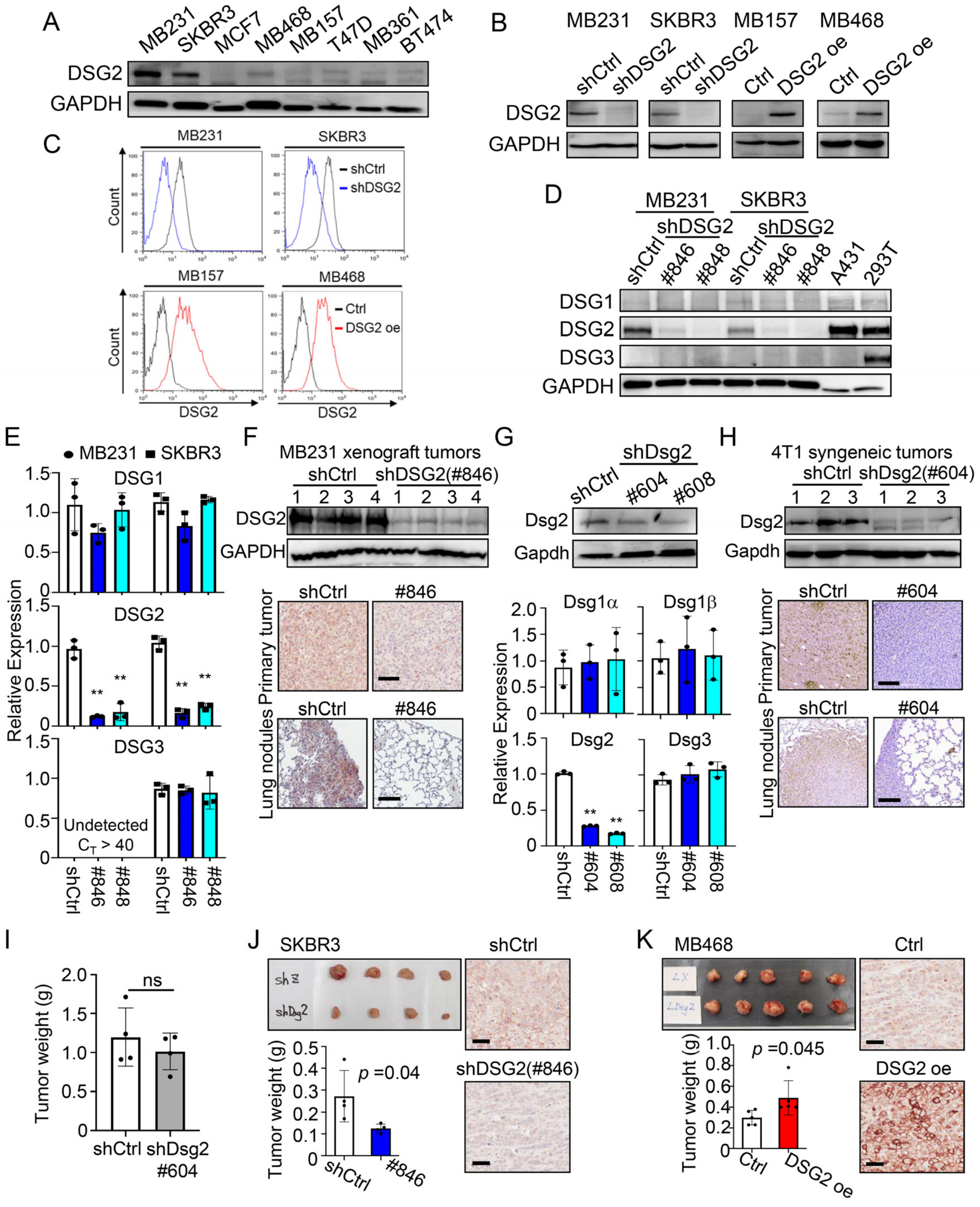


**Figure S2**

**(A)** IB of DSG2 expression in breast cancer cell lines.

**(B)** IB analyses of DSG2 expression in *DSG2*-depleted (shDSG2#846) MB231 and SKBR3 cells as well as *DSG2*-overexpressing MB157 and MB468 cells.

**(C)** FACS analyses of DSG2 expression in *DSG2*-depleted (shDSG2#846) MB231 and SKBR3 cells as well as *DSG2*-overexpressing MB157 and MB468 cells.

**(D)** IB of human DSGs in MB231 and SKBR3 cells transduced with shCtrl, shDSG2#846 or shDSG2#848. A431 and 293T cells were used as positive controls for DSG1 and DSG3, respectively. GAPDH is used as a loading control. Blots shown are from one representative experiment of three replicates.

**(E)** qRT-PCR of human DSGs in MB231 and SKBR3 cells transduced with shCtrl, shDSG2#846 or shDSG2#848. Three independent experiments were performed and data are means ± SD from one representative experiment (n =3). Significant differences are based on unpaired T-test. Average $C_T$ values for DSG genes in shCtrl cells: DSG1 (38.4), DSG2 (24.8) and DSG3 (40.1).

**(F)** Immunoblot of DSG2 expression in primary tumors and representative IHC staining of DSG2 in primary tumors and lung nodules of the MB231-EGFP shCtrl or shDSG2#846 tumor bearing mice. GAPDH is used as a loading control. Scale bar, 100 μm.

**(G)** Immunoblot (top) and qRT-PCR of mouse DSG2 in 4T1-GFP/LUC cells transduced with shCtrl, shDsg2#604 or shDSG2#608. Average $C_T$ values for Dsg genes in shCtrl cells: Dsg1α (40.2), Dsg1β (38.2), Dsg2 (27.3) and Dsg3 (31.9).

**(H)** Immunoblot of DSG2 expression in primary tumors and representative IHC staining of DSG2 in primary tumors and lung nodules of the 4T1-GFP/LUC shCtrl or shDSG2 tumor bearing mice. GAPDH is used as a loading control. Scale bar, 100 μm.

**(I)** Tumor weight of shCtrl or shDsg2#604 lentiviral vector transduced 4T1-GFP/LUC derived tumors from the mouse syngeneic model.

**(J)** Tumor weight and DSG2 IHC staining in SKBR3 derived tumors from xenograft models.

**(K)** Tumor weight and IHC staining in MB468 derived tumors from xenograph models. All data are presented as mean ± SD with significant differences detected by unpaired T-test.

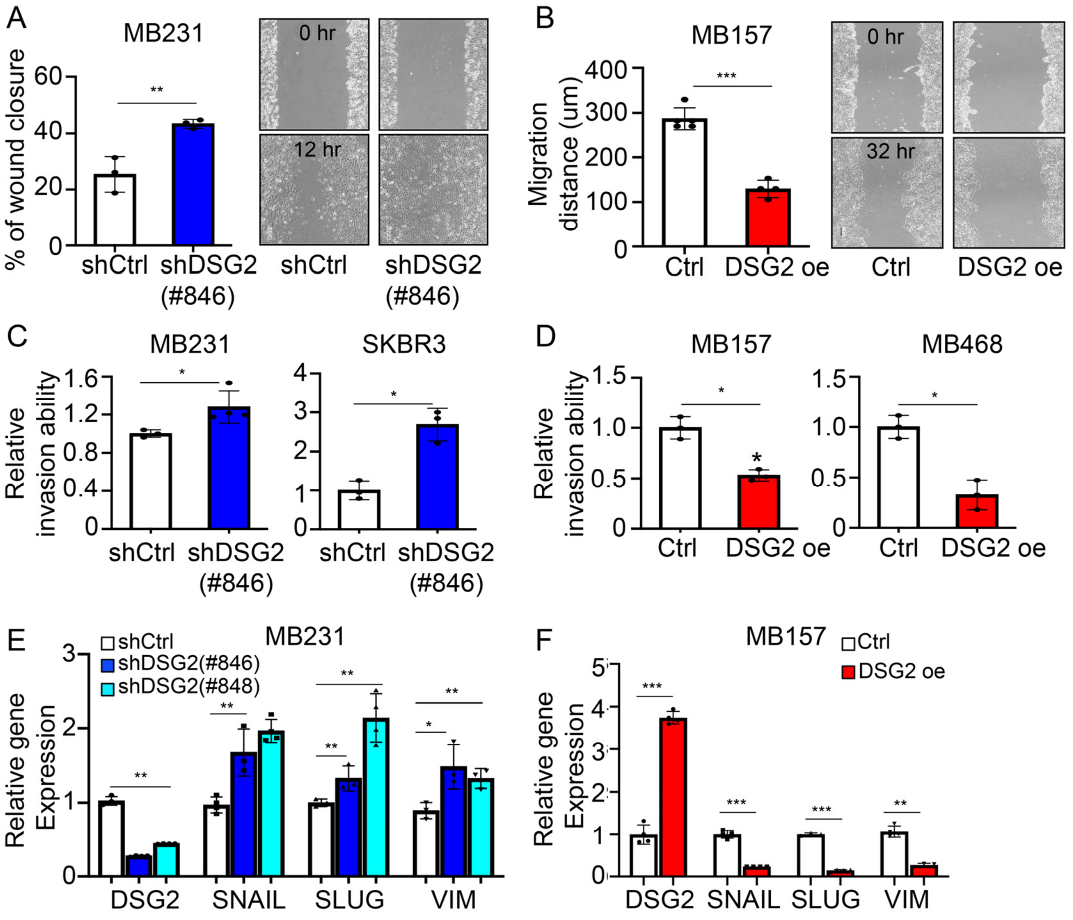


**Figure S3**

**(A and B)** Wound-healing migration assays using *DSG2*-depleted MB231cells (A) and *DSG2*-overexpressing MB157 cells (B).

**(C and D)** Transwell invasion assays using *DSG2*-depleted MB231 and SKBR3 cells (C), and *DSG2*-overexpressing MB157 and MB468 cells (D).

**(E, F)** qRT-PCR of *DSG2*, *SNAIL*, *SLUG* and *VIM* in *DSG2*-depleted SKBR3 cells (E) and *DSG2*-overexpressing MB157 cells (F). All data are presented as mean ± SD (significant differences detected by unpaired T-test). *** $p < 0.001$, **$p < 0.01$.

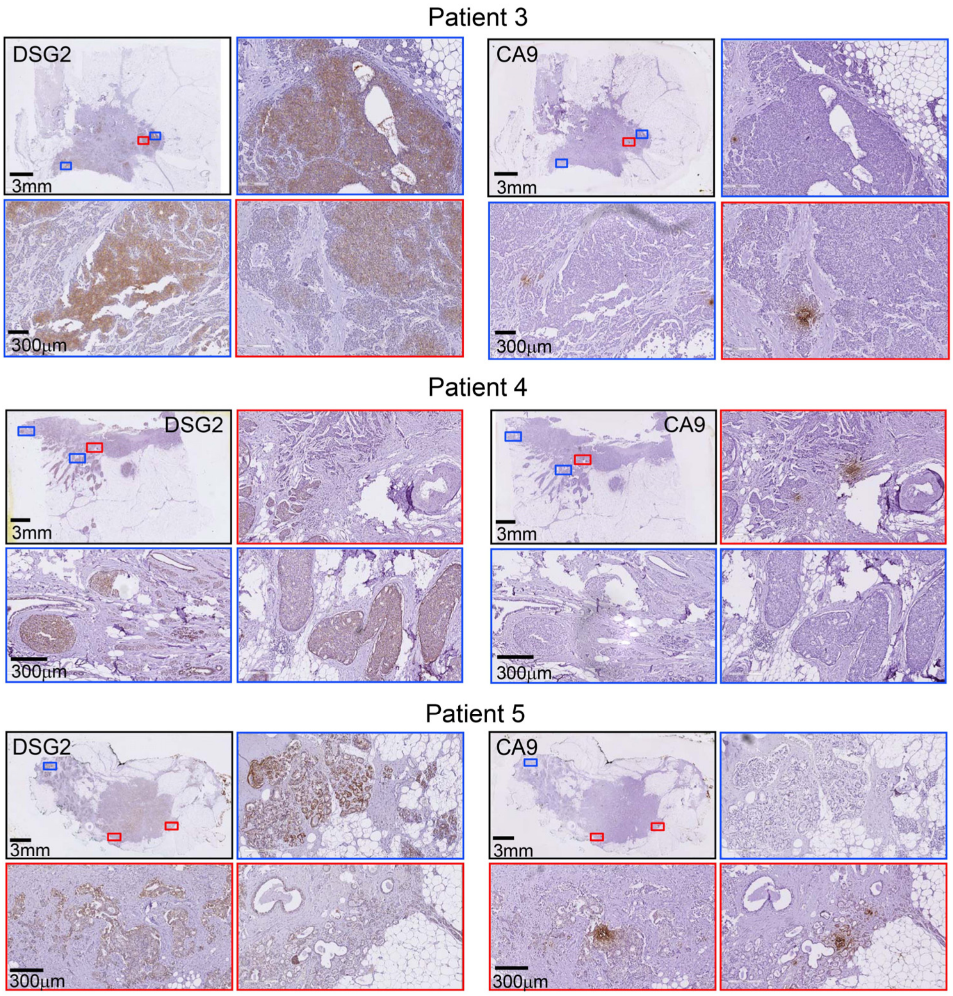


**Figure S4.**

Representative images of DSG2 and CA9 IHC staining using serial tumor sections from clinical breast cancer patients. Red boxes show enlarged images of low DSG2 but high CA9 expression regions. Blue boxes show enlarged images of high DSG2 but low CA9 expression regions. Scale bars indicate 3 mm or 300 µm as shown in the images. Examples shown are representative of 151 patient samples examined. Among the 151 samples, 66 slides contained $DSG2^{high}/CA9^{low}$ cancer cells only, 40 slides contained $DSG2^{low}/CA9^{low}$ cancer cells only and 45 slides contains both $DSG2^{high}/CA9^{low}$ and $DSG2^{low}/CA9^{high}$ cells in different regions.

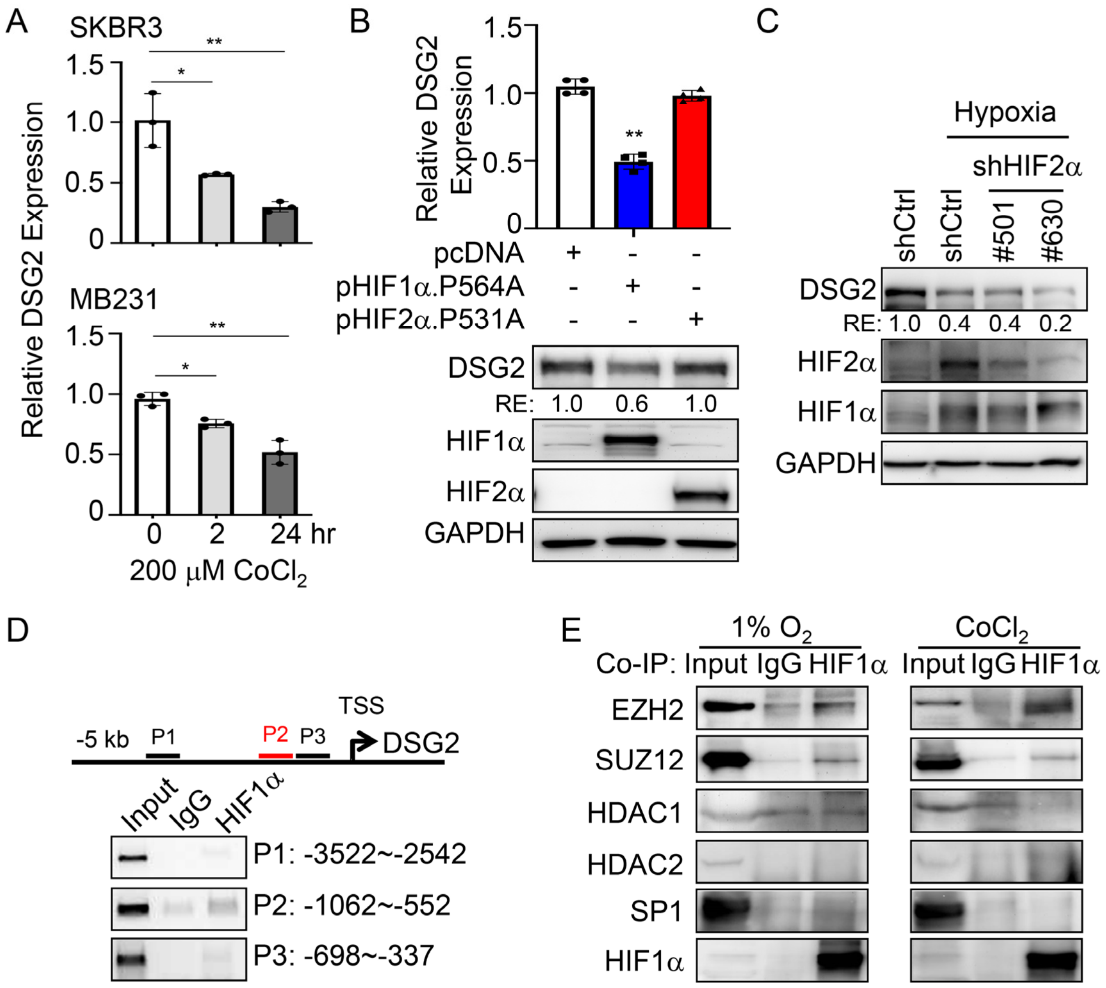


**Figure S5**

**(A)** qRT-PCR analysis of DSG2 in SKBR3 and MB231 cells treated with 200 μM CoCl2 for 2 or 24hr. Data are means ± S.D. (n =3). Significant differences are based on unpaired T-test. The experiments were repeated three times.

(B) qRT-PCR analysis of DSG2 level in 293T cells transiently transfected with pcDNA vector, pcDNA-HIF1α.P564A or pcDNA-HIF2α.P531A plasmid. Three independent experiments were performed and data are means ± SD from one representative experiment (n =3). Significant differences are based on unpaired T-test. DSG2, HIF1α and HIF2α expression was detected by immuoblot with GAPDH as a loading control.

Blots shown are from one representative experiment of three replicates. RE: relative expression.

(**C**) Immunoblot of DSG2 level in SKBR3 cells transduced with shHIF2α lentiviral vectors (clone # 501 and #630) and treated with hypoxia for 16 hours. DSG2, HIF1α and HIF2α expression was detected by immuoblot with GAPDH as a loading control. Blots shown are from one representative experiment of three replicates.

**(D)** Diagram shows the positions of three primer sets (P1 to P3 regions) flanking 5 kb upstream from the transcription start site (TSS) of the DSG2 promoter. These primers were used for ChIP–semiquantitative PCR analysis of HIF1α on the DSG2 promoter in SKBR3 cells under hypoxia for 8 hours.

**(E)** Co-IP of HIF1α, EZH2, SUZ12, HDAC1, HDAC2 and SP1 in SKBR3 cells under hypoxia or $CoCl_2$ treatment. IgG was used as a control. All data are presented as mean ± SD (significant differences detected by unpaired T-test). *** $p < 0.001$, **$p < 0.01$.

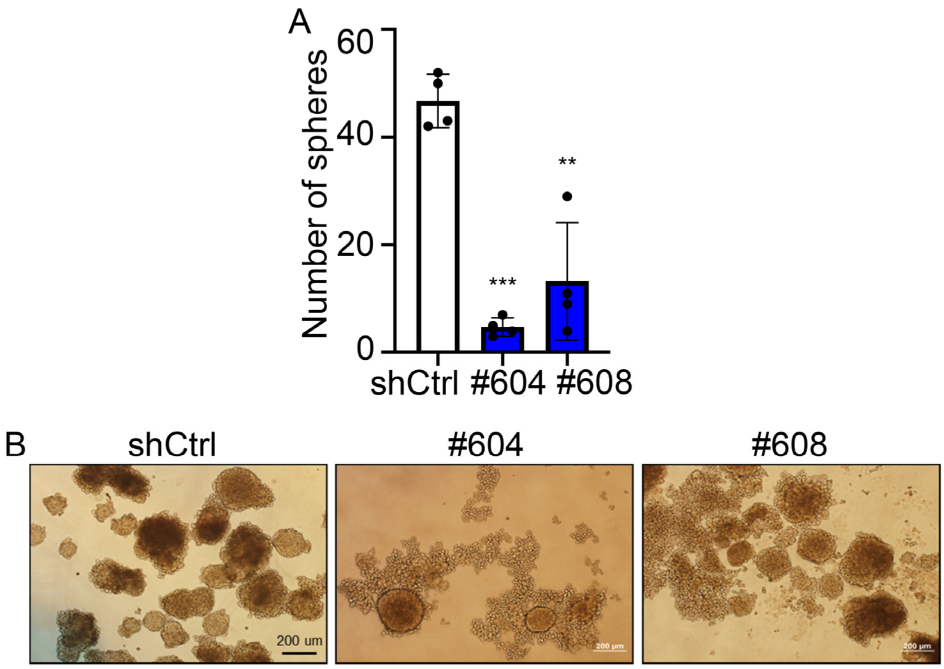


**Figure S6**

**(A)** Quantification of mammosphere forming ability using control (shCtrl) or *Dsg2*-depleted (#604 or #608) 4T1-GFP/LUC cells. All data are presented as mean ± SD (significant differences detected by unpaired T-test). **$p$ < 0.01, *** $p$ < 0.001.

**(B)** Representative images of mammospheres derived from the shCtrl- or shDsg2 (#604 or #608)-transduced 4T1-GFP/LUC cells in low-attached culture. Scale bar, 200 µm.

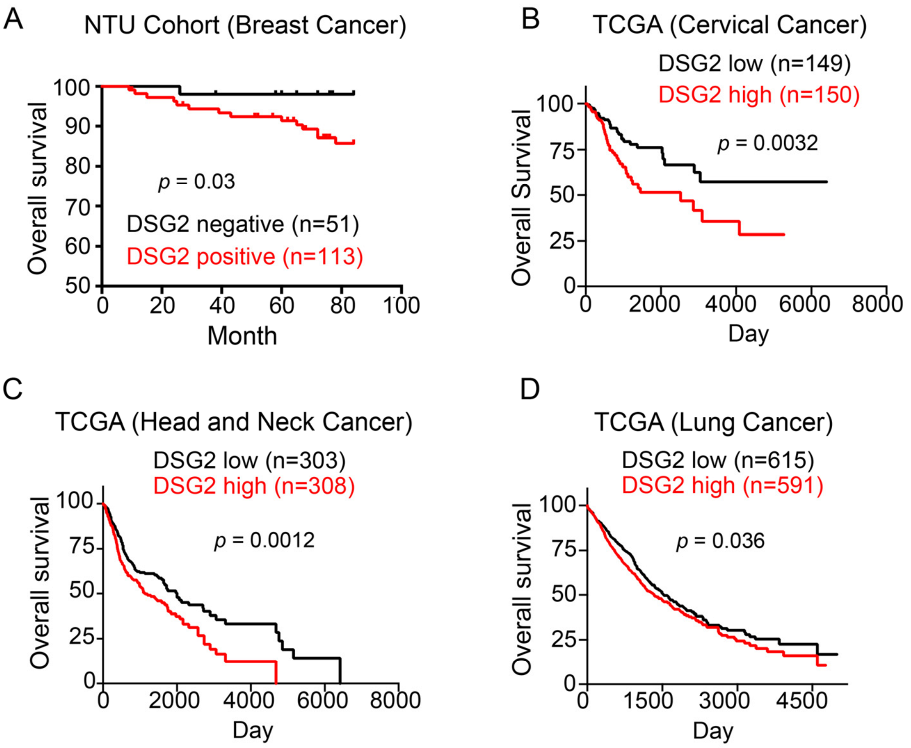


**Figure S7**

**(A)** Kaplan-Meier overall survival analysis of breast cancer patients grouped by DSG2 expression. The DSG2 positive group is indicated by red line (n = 113) and the DSG2 negative group is indicated by black line (n = 51). $p = 0.03$. The *p*-value was determined by log-rank test.

(**B-D**) Data from the Cancer Genome Atlas (TCGA) database analyzed by Kaplan-Meier overall survival analysis of cervical (B), head and neck (C) and lung (D) cancer patients grouped by *DSG2* expression. The results shown here are based upon data generated by the TCGA Research Network: https://www.cancer.gov/tcga. The mean *DSG2* expression level in each cancer was used as a cutoff value for classification of patients into *DSG2* high (DSG2 expression higher than the mean, indicated by red line) and *DSG2* low

(DSG2 expression lower than the mean, indicated by black line) groups. The *p*-value was determined by log-rank test.

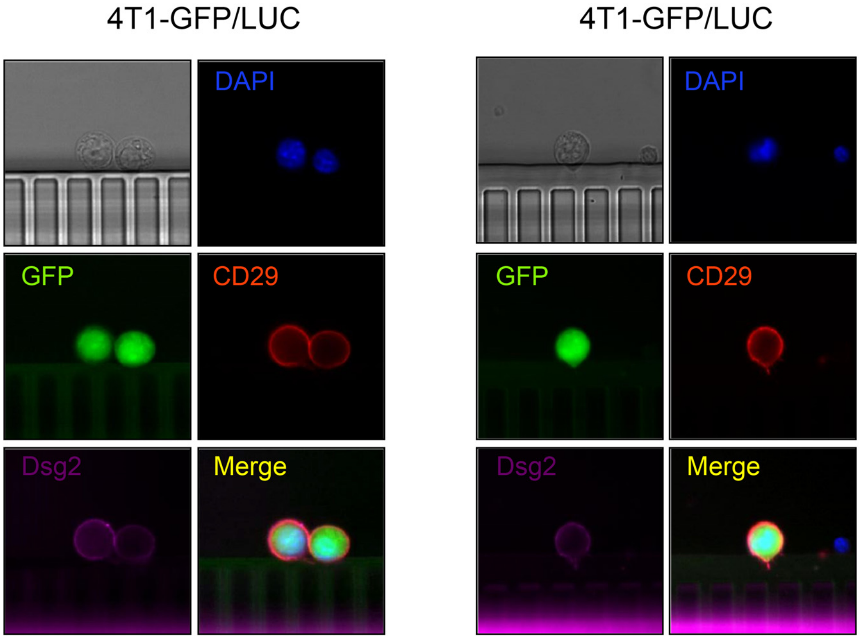


**Figure S8**

Spike-in control experiment to determine whether the centrifugation step in CTC analysis increases cell aggregation. The experiment used $10^3$ 4T1-GFP/LUC cells mixed with 2ml blood from wildtype BALB/c mice. 4T1-GFP/LUC cells were identified using GFP, CD29 and DAPI. No CTC clusters were detected.

**Appendix Tables**

**Table S1. Primer sequence**

| **Gene** | **Forward (5'- 3')** | **Reverse (5'- 3')** |
|---|---|---|
| **Primers for qRT-PCR** | | |
| DSG1 | CTTCTGAACCCGGAAACGGA | ATGAGGAGTCCAATGCCAGC |
| DSG2 | TGGATCAAGGGGGCAGTCTA | TCAGTGGTCATGATAGCGCC |
| DSG3 | TGTAGACTCCTTCGGAAAGC A | TAGTTCTGGGGAAGAGCCCC |
| Dsg1α | AATACCAAAGGCTTAATGGG GAA | CTGGAACTGCCACCACTTTCT |
| Dsg1β | AATACCAAAGGCTTAATGGG GAAT | CACCACCATCTGAACCTGGTA |
| Dsg2 | CGTACTCCTCTAACACCGGC | GGTGGCGGTACTACCTTCTG |
| Dsg3 | CTGTGACTGTGGGTCAGGTC | CTGAGCTCCTTCGATTCCCC |
| SNAIL | CCTCCCTGTCAGATGAGGAC | CCAGGCTGAGGTATTCCTTG |
| SLUG | GGGGAGAAGCCTTTTTCTTG | TCCTCATGTTTGTGCAGGAG |
| VIM | CAGATGCGTGAAATGGAAG A | TGGAAGAGGCAGAGAAATCC |
| GAPDH | GGCTCTCCAGAACATCATCC CTGC | GGGTGTCGCTGTTGAAGTCAG AGG |
| **Primers for ChIP** | | |
| P1 | GATGGGACCCGATGCTAGA A | GTTGTGGTCCCATTGGTAGC |
| P2 | TCTCACCTCGCAATCACGTT | AAAGGTCCCGCACTTTGTGA |
| P3 | CTGTTGGAAGCAGTGAAGG G | GTTCTCTCCTTCCCGACCC |
| ChIP-qPCR | CAAAACTTCCCCGTGCGTTG | AAAGGTCCCGCACTTTGTGA |

**Table S2. Antibodies**

| **Gene** | **Company (Cat. #)** | **Dilution** |
|---|---|---|
| **Antibodies for Immunoblotting/IHC** | | |
| DSG1 | Genetex (Cat.129983) | 1:1000 |
| DSG2 | homemade | 1:1000/1:300 |
| DSG3 | Genetex (Cat.129931) | 1:1000 |
| mouse DSG2 | Novus (Cat. 75469) | 1:1000/1:300 |
| HIF1α | Genetex (Cat.127309) | 1:1000 |
| HIF1α | BD (Cat.610958) | 1:1000 |
| HIF2α | Genetex (Cat.632015) | 1:1000 |
| SUZ12 | Cell Signaling (Cat.3737) | 1:1000 |
| EZH2 | Cell Signaling (Cat.5246) | 1:1000 |
| HDAC1 | Cell Signaling (Cat.5356) | 1:1000 |
| SP1 | Cell Signaling (Cat.5931) | 1:1000 |
| EGFR | Cell Signaling (Cat.4267) | 1:1000 |
| HDAC2 | Cell Signaling (Cat.25451) | 1:1000 |
| GAPDH | Genetex (Cat. GT239) | 1:10000 |
| CAIX | Genetex (Cat.70020) | 1:500 |
| Anti-rabbit HRP | Bethyl (Cat.A120-12P) | 1:10000 |
| Anti-mouse HRP | Bethyl (Cat.A90-516D2) | 1:10000 |
| **Antibodies for CoIP (for 250 μg total protein) and ChIP (for 200 μg chromatin)** | | |
| HIF1α | Genetex (Cat.127309) | 4 ug |
| SUZ12 | Cell Signaling (Cat.3737) | 4 ug |
| EZH2 | Cell Signaling (Cat.4905) | 4 ug |
| H3K27me3 | Cell Signaling (Cat.9756) | 2 ug |
| **Antibodies for Immunofluorescence or FACS** | | |
| Anti-mouse 488 | Invitrogen (Cat.A28175) | 1:400 |
| Anti-mouse 647 | Sigma-Aldrich (Cat.SAB4600182) | 1:500 |
